# Single nuclei sequencing reveals C_4_ photosynthesis is based on rewiring of ancestral cell identity networks

**DOI:** 10.1101/2023.10.26.562893

**Authors:** Joseph Swift, Leonie H. Luginbuehl, Tina B. Schreier, Ruth M. Donald, Travis A. Lee, Joseph R. Nery, Joseph R. Ecker, Julian M. Hibberd

## Abstract

In multicellular systems changes to the patterning of gene expression drive modifications in cell function and trait evolution. One striking example is found in more than sixty plant lineages where compartmentation of photosynthesis between cell types allowed the evolution of the efficient C_4_ pathway from the ancestral C_3_ state. The molecular events enabling this transition are unclear. We used single nuclei sequencing to generate atlases for C_3_ rice and C_4_ sorghum during photomorphogenesis. Our analysis revealed that initiation of photosynthesis gene expression is conditioned by cell identity. In both species a conserved cistrome defines each cell type, and photosynthesis genes switching expression from mesophyll in rice to bundle sheath in sorghum acquire hallmarks of bundle sheath identity. The sorghum bundle sheath has also acquired gene networks associated with C_3_ guard cells. We conclude C_4_ photosynthesis is based on rewiring in *cis* that exapts cell identity networks of C_3_ plants.

## Introduction

Multicellularity has evolved repeatedly such that it is now found in multiple lineages of bacteria and fungi, as well as the metazoans, algae and land plants (Grosberg and Strathmann, 2007). In all cases it allows particular cells to become specialized to carry out specific functions. In land plants, the division of labor between different cell types underlies many important agricultural traits such as water and nutrient uptake in roots and photosynthesis in shoots. In the case of photosynthesis, for the majority of land plants, CO_2_ fixation occurs primarily in mesophyll cells and is dependent on the enzyme Ribulose-1,5-Bisphosphate Carboxylase-Oxygenase (RuBisCO). Since the first fixation product of RuBisCO is the three-carbon metabolite 3-phosphogylceric acid, this pathway has been termed C_3_ photosynthesis (Bassham, Benson, Calvin 1950). It is found in bacteria and algae as well as land plants and so is considered ancestral. In land plants, although mesophyll cells are fundamental for photosynthesis, the specialization of other cell types is equally important (Haberlandt, 1884). For example, the xylem transports water and nutrients, guard cells mediate the flow of water and CO_2_ in and out of the leaf, while the epidermis protects from the external environment. Similarly, once sugars are synthesized in the mesophyll, the bundle sheath cells deliver them to the phloem for allocation to other tissues.

Although most land plants use the C_3_ pathway, RuBisCO is not able to completely discriminate between CO_2_ and O_2_. In addition to loss of carbon fixation, when RuBisCO carries out oxygenation reactions it generates the toxic intermediate phosphoglycolate that needs to be rapidly metabolized via the energy-intensive photorespiratory cycle (Bowes et al., 1971). At higher temperatures the proportion of oxygenation reactions increases, and so the efficiency of C_3_ photosynthesis declines (Bauwe et al., 2010). In multiple plant lineages, including staple crops such as maize and sorghum, evolution has reconfigured the functions of mesophyll and bundle sheath cells such that CO_2_ fixation by RuBisCO is repressed in the mesophyll and activated in the bundle sheath. These species are known as C_4_ plants because the first committed step of the pathway requires substantial activities of phospho*enol*pyruvate carboxylase in mesophyll cells and produces the C_4_ acid oxaloacetate. Reduction or transamination of oxaloacetate to malate and aspartate in the mesophyll allows C_4_ acids to accumulate to high concentrations and diffuse into bundle sheath cells prior to decarboxylation in close proximity to RuBisCO (Hatch and Slack, 1966). This process leads to a tenfold increase in CO_2_ concentration in bundle sheath chloroplasts (Furbank, 2011), which reduces the frequency of oxygenation reactions so that photosynthetic as well as water and nitrogen use efficiencies are optimized (Ghannoum et al., 2010). As a result, C_4_ plants grow particularly well in hot and dry climates and constitute some of the most productive crop species in the world (Jordan and Ogren, 1984; Sage and Zhu., 2011). Crucially, in both C_3_ and C_4_ plants, photosynthetic efficiency is dependent on mechanisms that pattern and maintain differential gene expression between each cell type of the leaf. However, it is not clear how strict partitioning of photosynthesis between cells is established and maintained, nor how these patterns change to allow the evolution of C_4_ photosynthesis from the ancestral C_3_ pathway.

In eukaryotes, the regulation of gene expression in distinct cell types and in response to endogenous and environmental signals is encoded by *cis*-regulatory elements, the binding sites of transcription factors (Marand et al., 2017). In C_4_ leaves mechanical and cell digestion approaches have provided insight into the patterning of gene expression in mesophyll and bundle sheath strands (Markelz et al., 2003; Covshoff et al., 2013; John et al., 2014; Burgess et al., 2019; Borba et al., 2023) and a small number of *cis*-regulatory mechanisms controlling cell specific expression of C_4_ genes have been identified (Gowik et al., 2004; Brown et al., 2011; Williams et al., 2016; Reyna-Llorens et al., 2018; Burgess et al., 2019). However, in almost all studies of C_4_ leaves, only bundle sheath strands have been examined, which contain phloem and xylem cells as well as the bundle sheath (Burgess et al., 2019, Borba et al., 2023). Separation of bundle sheath cells from C_3_ leaves has been even more challenging (Aubry et al., 2014) and so molecular mechanisms allowing the rewiring of gene expression during the evolution from C_3_ to C_4_ photosynthesis are poorly understood. We reasoned that the advent of single nuclei sequencing (Grindberg et al., 2013; Tian et al., 2020; Cervantes-Perez et al., 2022; Guillotin et al., 2023; Lee et al., 2023; Nobori et al., 2023; Sun et al., 2023; Wang et al., 2023) may remove this substantial roadblock by allowing the transcriptional identity of each cell type in C_3_ and C_4_ species to be defined. To achieve this, we selected rice and sorghum that use the C_3_ and C_4_ pathways respectively to generate single nuclei atlases for gene expression and chromatin accessibility. Both species are models and diploid crops of global importance representing distinct clades in the monocotyledons that diverged approximately 81 million years ago (Huang et al., 2022). Molecular signatures of each cell type that are shared by rice and sorghum may therefore also be found in these cells in the ∼11,000 species derived from their last common ancestor.

To define how patterning of photosynthesis gene expression is established we sampled nuclei after transfer of seedlings from dark to light, a stimulus that induces photomorphogenesis and thus activation of photosynthesis gene expression. In both species gene expression was rapidly induced by light in all cell types. However, our data show that prior to the perception of light, the expression and chromatin accessibility of many photosynthesis genes is conditioned by cell identity. Thus, cell identity defines the extent to which cell types can respond to light. Furthermore, we found that between species, changes in transcriptional cell identity are the most dramatic in the bundle sheath, allowing the C_4_ bundle sheath to change its response to light. This is driven in part by the C_4_ bundle sheath gaining cell identity markers from mesophyll and guard cells of C_3_ leaves. While transcriptional cell identities can change across species, we found that the underlying *cis*-elements that define cell identity are conserved. Genes that rewire their expression to become bundle sheath specific in C_4_ sorghum acquire ancestral *cis*-elements such as DNA-binding with One Finger (DOF) motifs that direct bundle sheath expression in both the C_3_ and C_4_ bundle sheath. The simplest explanation from these findings is a model in which the evolution of photosynthesis is based on C_4_ genes acquiring *cis*-elements associated with bundle sheath identity that then harness a stable patterning of transcription factors between cell types of C_3_ and C_4_ leaves.

## Results

### Single-nuclei gene expression and chromatin accessibility atlases for rice and sorghum shoots during de-etiolation

To understand how different cell types in rice and sorghum shoots respond to light, we grew seedlings of each species in the dark for 5 days and then exposed them to a light-dark photoperiod for 48 hours (**Figure 1A**). As expected, shoot tissue underwent photomorphogenesis during this time. For example, leaves emerged from the shoot and chlorophyll accumulated within the first 12h of de-etiolation (**Figure 1A, Figure S1**). Scanning Electron Microscopy (SEM) showed that leaves uncurled in response to light (**Figure 1B**) and that etioplasts in both mesophyll and bundle sheath cells contained prolamellar bodies before light exposure (**Figure 1C**). Within 12h of light etioplasts had converted into mature chloroplasts with assembled thylakoid membranes. Compared with rice, chloroplast development was more pronounced in the bundle sheath of sorghum, and clear differences in thylakoid stacking in chloroplasts from mesophyll and bundle sheath of sorghum were evident (**Figure 1C**).

**Figure 1:**
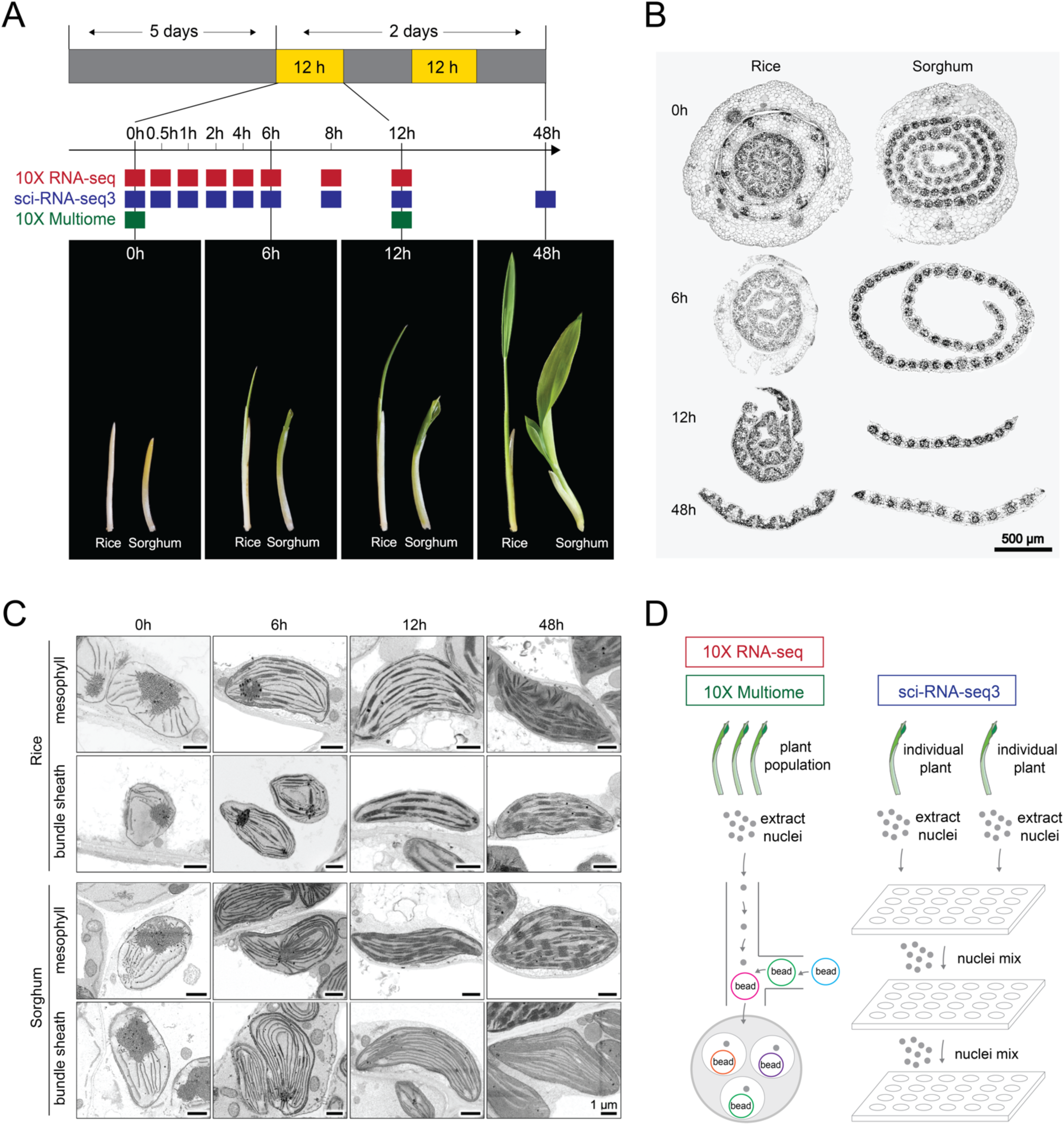
Photomorphogenesis of rice and sorghum shoots during de-etiolation. **(A)** Schematic of de-etiolation time course. Plants were grown in the dark for 5 days before exposure to light for 2 days. **(B)** Scanning electron micrographs of rice and sorghum leaf cross sections showing leaf maturation from 0h to 48h after light exposure. **(C)** Scanning electron micrographs of etioplasts and chloroplasts from mesophyll and bundle sheath cells of rice and sorghum shoots at 0h, 6h, 12h, and 48h after light exposure. **(D)** Summary of 10X Genomics platform used for RNA and ATAC-sequencing of single nuclei extracted from a population of plants, and sci-RNA-seq3 used for sequencing single nuclei from individual plants to provide increased biological replication.

Underlying this cellular remodeling and activation of the photosynthetic apparatus during photomorphogenesis are changes in gene regulation. However, to date these have been described in bulk tissue samples and so how each cell type responds is not known (Boffey et al., 1980; Sullivan et al., 2014; Armarego-Marriott et al., 2019; Armarego-Marriott et al., 2020; Singh et al., 2023). To better understand how cells respond to light we generated single nuclei atlases of transcript abundance for both rice and sorghum shoots as they undergo photomorphogenesis. To achieve this, shoot tissue at nine time points during de-etiolation was harvested and nuclei sequenced using 10X and sci-RNA-seq3 (Cao et al., 2019) (**Figure 1A, 1D**). By combining nuclei sampled over the time course from both sequencing approaches we generated gene expression atlases derived from 190,569 and 265,701 nuclei from rice and sorghum respectively. Using the 10X-multiome workflow (combining RNA-seq and ATAC-seq) we also assayed cell specific changes in chromatin accessibility at 0h and 12h after light exposure by sequencing 22,154 and 20,169 nuclei from rice and sorghum respectively.

These outputs were visualized using uniform manifold approximation projection (UMAP) and nineteen distinct clusters were identified for each species (**Figure 2A, 2B, Figure S2**). Using the expression of previously described marker genes and their orthologs, cell types were assigned to each main cluster. This included mesophyll, guard, epidermal, xylem parenchyma and phloem cells (including parenchyma and companion cells) (**Figure 2C, 2D**). Gene Ontology (GO) terms derived from cluster-specific genes reflected previously documented functions for each cell type (**Table S1**). For example, mesophyll nuclei showed high expression of genes involved in photosynthesis, and clusters containing nuclei from epidermis cells were enriched in genes involved in lipid biosynthesis and export, consistent with the role of this tissue in cutin production (Suh et al., 2005). Moreover, nuclei from phloem and xylem primarily expressed genes for transport of water and solutes and the synthesis of cell wall components respectively (**Table S1**). Each cluster contained nuclei sampled from all time points, indicating that clustering was driven predominantly by cell type rather than time after exposure to light (**Figure S2**).

**Figure 2:**
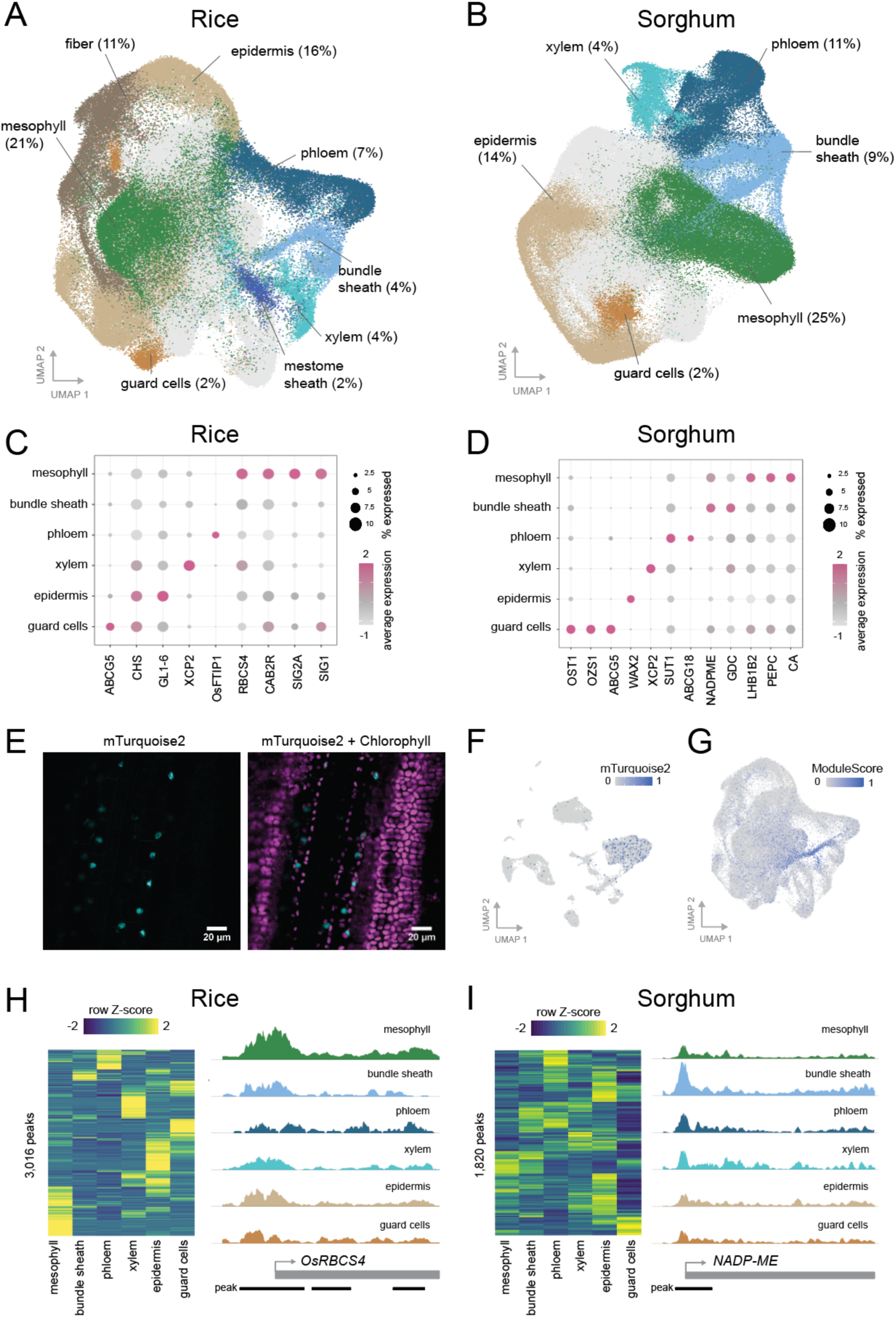
Single cell atlases for gene expression and chromatin accessibility in rice and sorghum shoots during de-etiolation. UMAP of transcript profiles of **(A)** rice and (**B)** sorghum single nuclei, encompassing all time points. Transcript abundance from marker genes in cell types of **(C)** rice and **(D)** sorghum. **(E)** Image from confocal laser scanning microscopy of a rice bundle sheath marker line expressing mTurquoise2 driven by the bundle sheath specific Zj*PCK* promoter. **(F)** Clustering of sequenced nuclei sourced from the mTurquoise2 rice bundle sheath marker line - whole leaf nuclei were enriched with mTurquoise2 nuclei before sequencing. **(G)** Expression of bundle sheath markers from **(F)** in the rice shoot de-etiolation single nuclei dataset from **(A). (H)** 3,016 peaks in accessible chromatin could be assigned to specific cell types in rice nuclei. Accessibility for the promoter of *OsRBCS4* in each cell type shown to the right. **(I)** 1,820 peaks in accessible chromatin could be assigned to specific cell types in sorghum nuclei. Accessibility of promoter region for *NADP-ME* in each cell type shown to the right.

We identified cells of the sorghum bundle sheath through expression of C_4_ cycle genes such as NADP-Malic Enzyme (*NADP-ME*) and Glycine Decarboxylase (*GDC*) (**Figure 2D**). However, to our knowledge there are no such markers for the bundle sheath in rice undergoing photomorphogenesis. To address this, we generated a stable reporter line in which bundle sheath nuclei were labelled with a fluorescent mTurquoise2 reporter under control of the *PHOSPHOENOLCARBOXYKINASE* promoter from *Zoysia japonica* (Nomura et al., 2005). Confocal laser scanning microscopy of these plants confirmed the presence of mTurquoise2 specifically in nuclei of rice bundle sheath cells (**Figure 2E**). A preparation of nuclei from whole leaves was enriched in bundle sheath nuclei obtained after fluorescence activated sorting of nuclei from this reporter line. Sequencing and clustering produced fourteen clusters, with the largest specifically expressing mTurquoise2, thus identifying nuclei from the rice bundle sheath (**Figure 2F, Figure S3**). Within this cluster we detected specific expression of genes such as *PLASMA MEMBRANE INTRINSIC PROTEIN (PIP1.1)*, *SULFITE REDUCTASE* (*SIR)* and *ATP SULFURYLASE (ATPSb)*, which have previously been shown by laser capture microdissection to be expressed in the bundle sheath from mature rice leaves (Hua et al., 2021) (**Table S2**). Using marker genes from this cluster it was then possible to annotate nuclei with bundle sheath identity in our de-etiolation dataset (**Figure 2F, 2G, Figure S3**).

Complementing this atlas describing cell-type gene expression, the multiome assay (RNA-seq and ATAC-seq) allowed changes in chromatin accessibility during photomorphogenesis to be detected. After cross-validation with the single nuclei transcriptional atlases, the multiome atlases identified six cell types from each species (**Figure S4, S5**). Between 1,820 and 3,016 accessible peaks in promoter regions were specific to each cell type (**Figure 2H, 2I**). As would be expected, these peaks were upstream of genes associated with the GO terms enriched in each cell type (**Table S3**). Peaks were also detected upstream of canonical marker genes for each cell type. For example, the promoters of RuBisCO small subunit (*RbcS4)* from rice and *NADP-ME* from sorghum were most accessible in mesophyll and bundle sheath cells respectively (**Figure 2H, 2I**).

### The C_4_ bundle sheath acquires a new transcriptional identity from C_3_ mesophyll and guard cells

C_4_ evolution has repeatedly repurposed the bundle sheath to perform photosynthesis (Hibberd and Covshoff 2010, Langdale 2011). However, because it has not previously been possible to define gene expression in bundle sheath cells from C_3_ or C_4_ plants, the extent to which this cell type has been altered is not known. Using our single nuclei atlas from rice and sorghum we therefore tested whether transcriptional rewiring of C_4_ bundle sheath cells is driven only by acquisition of photosynthesis networks associated with mesophyll cells of C_3_ plants, or whether more substantial transcriptional changes are involved. To understand how the transcriptional identities of each cell type from rice and sorghum differ, we generated a pan-transcriptome atlas of photosynthetic tissue sampled at 48h after light exposure. Despite the evolutionary distance between rice and sorghum, most cell types from these species co-clustered (**Figure 3A, Figure S6**). In contrast, nuclei from bundle sheath cells in rice and sorghum did not co-cluster (**Figure 3A**). Supporting this observation, GO enrichment analysis indicated that cells of the C_3_ and C_4_ bundle sheath carry out distinct functions - while genes expressed in the bundle sheath of rice were predominantly associated with transport and localization, those of sorghum were associated with organic acid metabolism and generation of precursor metabolites and energy (**Table S1**).

**Figure 3:**
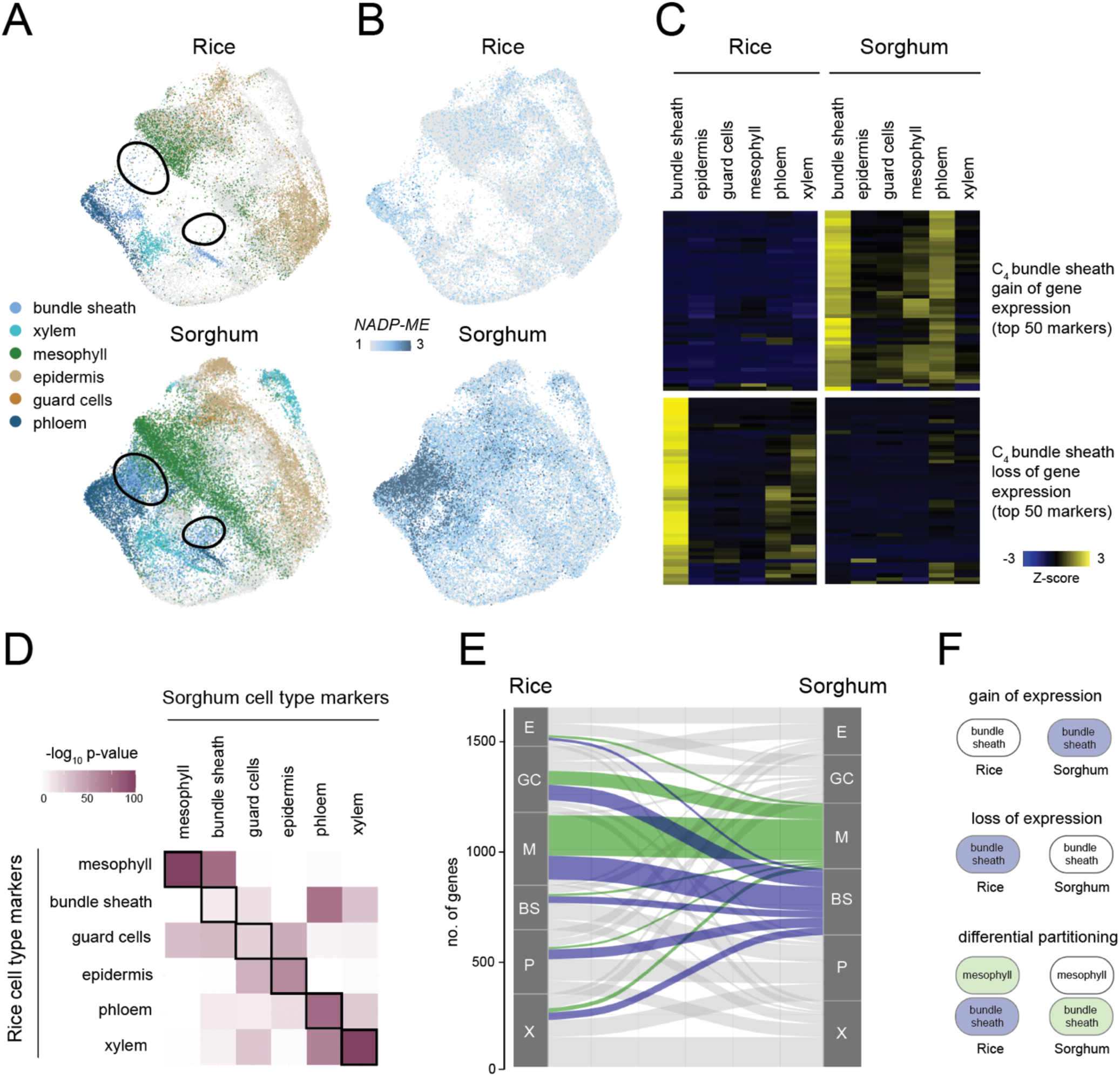
The C_4_ sorghum bundle sheath derives a transcriptional identity associated with mesophyll and guard cells found in C_3_ rice. **(A)** Pan-transcriptome UMAP for rice and sorghum nuclei 48h after light exposure. UMAPs (Uniform Manifold Approximation and Projection) indicate rice nuclei (top) and sorghum nuclei (bottom). Areas indicated with a black circle show bundle sheath nuclei from sorghum. **(B)** Transcript abundance for sorghum *NADP-ME* and its rice ortholog at 48h after light exposure. **(C)** Heatmap of transcript abundance for bundle sheath marker genes in rice and sorghum in each cell type 48h after exposure to light. **(D)** Statistical significance of overlap between cell type specific marker genes of rice and sorghum. **(E)** Sankey plot summarizing changes in partitioning of marker genes between cell types. Marker genes for sorghum mesophyll and bundle sheath cells indicated in green and blue respectively. **(F)** Schematic illustrating three types of gene expression changes that underpin re-functionalization of the C_4_ bundle sheath.

More than 180 genes that were specific to bundle sheath cells of sorghum had orthologs that were either poorly expressed or were not specific to any cell type in rice (**Table S4**). For example, the canonical C_4_ gene *NADP-ME* was strongly and specifically expressed in the sorghum bundle sheath but was poorly expressed in rice in a non cell type specific manner (**Figure 3B**). Similar patterns of high and localized expression were evident in the sorghum bundle sheath for other genes involved in photosynthesis, photorespiration and chloroplast functions (**Figure 3C, Table S4**). The bundle sheath of C_4_ sorghum also lost expression of genes associated with this cell type in rice (**Figure 3C, Table S4**). Interestingly, this included genes involved in hormone signaling and biosynthesis including gibberellic acid, ethylene, and auxin pathways, as well as genes encoding sugar and water transporters. We next investigated how conserved cell type specific gene expression patterns were across species. While most cell types showed conserved patterns of expression between rice and sorghum, this was not the case for the bundle sheath (**Figure 3D**). In fact, transcripts from only 28 orthologs (including genes involved in sulfur metabolism and transport) were specific to the bundle sheath of both species (**Figure 3D, 3E, Table S4**). The C_4_ bundle sheath of sorghum had also obtained patterns of gene expression from other cell types (**Figure 3E**). Indeed, bundle sheath cells of sorghum were transcriptionally more similar to mesophyll and guard cells of rice, whereas the bundle sheath of rice was most similar to the phloem of sorghum (**Figure 3D**, **Figure 3E**). Similarities between sorghum bundle sheath and rice mesophyll or guard cells were primarily driven by changes in the expression of genes involved in the Calvin-Benson-Bassham Cycle and starch metabolism (**Table S5**). Thus, to transcriptionally rewire the C_4_ bundle sheath it appears that this cell type (***i***) gained genes not found specifically or highly expressed in rice shoot tissue, (***ii***) lost specific expression of genes transcribed in the C_3_ bundle sheath, and (***iii***) gained genes preferentially expressed in other cell types of C_3_ rice including mesophyll and guard cells (**Figure 3F**).

As a difference in expression of photosynthesis genes between bundle sheath and mesophyll cells (hereafter partitioning) is considered crucial for the evolution of C_4_ photosynthesis (Hibberd and Covshoff 2010) we examined this phenomenon in sorghum and rice. Pairwise comparison of gene expression in response to light revealed that in each species transcripts from more than one thousand genes were partitioned between mesophyll and bundle sheath cells and included 225 orthologous gene pairs (**Figure 4A, Table S6**). Of these, 126 were partitioned identically between the same cell types in both rice and sorghum (**Figure 4B**). Those consistently partitioned to the mesophyll in both species were associated with oxidation and reduction processes as well as carbohydrate and small molecule metabolism, while bundle sheath partitioned genes were involved in transport of solutes (**Figure 4B, Table S6**). Interestingly, an additional 99 orthologs showed opposing patterns in the two species, i.e. they were ‘differentially’ partitioned. 43 orthologs that had swapped from strong expression in the mesophyll of rice to strong expression in the bundle sheath of sorghum included genes encoding proteins of the Calvin-Benson-Bassham cycle as well as organic acid and nitrogen metabolism. 56 genes that swapped from strong expression in the bundle sheath of rice to the mesophyll of sorghum were associated with transport of metabolites and solutes (**Figure 4B**).

**Figure 4:**
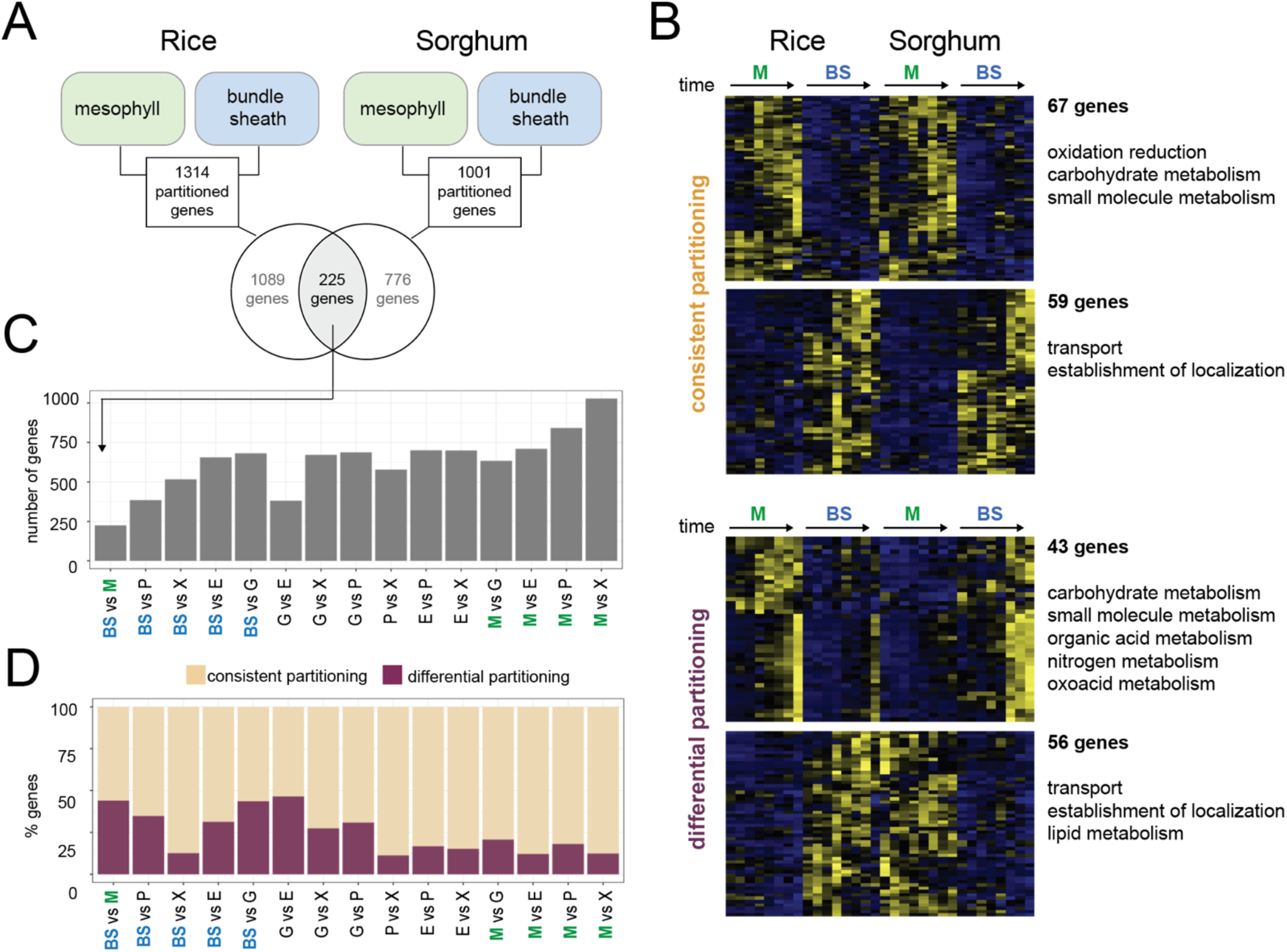
Bundle sheath cells show the least conservation in transcript partitioning between rice and sorghum. **(A)** Number and overlap of genes differentially expressed between mesophyll and bundle sheath cells in rice and sorghum. **(B)** Differentially expressed orthologs between mesophyll and bundle sheath cells of rice and sorghum. Genes fall into two categories, those consistently partitioned, i.e. more highly expressed in the same cell type in both rice and sorghum, and those that are differentially partitioned, i.e. swap expression from one cell type to the other. Gene ontology terms associated with genes that fall into each category are shown on the right. **(C)** Quantifying overlap of genes partitioned between different cell type pairs in rice and sorghum. **(D)** Percentage of partitioned genes in **(C)** that are either differentially or consistently partitioned.

To investigate how conserved partitioning was between all cell types, we assessed the degree of cross-species overlap between each pair of the six cell types annotated (**Figure 4C**). This revealed that the mesophyll and bundle sheath had the smallest set of partitioned genes across species, as well as the weakest statistical overlap (**Figure 4C**, **Figure S7**). In addition to mesophyll and bundle sheath cells of rice showing the lowest conservation in terms of transcript partitioning, it was also noticeable that a large proportion of the genes that were partitioned between these cells had in fact swapped cell types (**Figure 4D**). This suggests that the swapping of functions or ‘identity’ between other cell types is a rare event genome-wide but occurs relatively frequently between the mesophyll and bundle sheath.

### Pervasive acquisition of light regulation within the bundle sheath of sorghum

Since light induces photomorphogenesis we next investigated how individual nuclei from each cell type responded to this stimulus. When rice mesophyll and sorghum bundle sheath nuclei were analysed they naively clustered by time of sampling, indicating that light was a dominant driver of transcriptional state (**Figure 5A, 5B**). Canonical marker genes showed the expected induction, for example, *RbcS* and *NADP-ME* were activated by light in mesophyll cells of rice (**Figure 5A**) and bundle sheath cells of sorghum (**Figure 5B**) respectively. We detected global cell-type specific differential gene expression responses to light by fitting statistical models to pseudo-bulked transcriptional profiles. Then, to find dominant expression trends in gene regulation, we clustered differentially expressed genes using Pearson correlation. In rice each of the six cell types examined showed a distinct and cell type specific response to light (**Figure 5C**, **Figure S8**, **Table S7**). Apart from the bundle sheath and epidermal cells in rice, hundreds of cell type-specific light-responsive genes were detected (**Figure 5C**). Notably, while genes in rice mesophyll cells showed a steady increase in expression, those in other cells showed a more complex response with multiple phases (**Figure 5C & 5D**). In both species, mesophyll and bundle sheath specific genes were enriched in photosynthesis and chloroplast-related functions, consistent with the rapid greening of shoots and the conversion of etioplasts into chloroplasts during de-etiolation (**Table S7**). Bundle sheath cells from rice and sorghum showed the greatest difference in their response to light. This change in behavior appears to be due to at least two phenomena. First, some genes expressed in bundle sheath cells of sorghum showed the same response to light as those expressed in the mesophyll (**Figure 5D**). Second, some transcripts partitioned to bundle sheath cells of sorghum showed a strong increase in abundance from 6 hours of light (**Figure 5D**).

**Figure 5:**
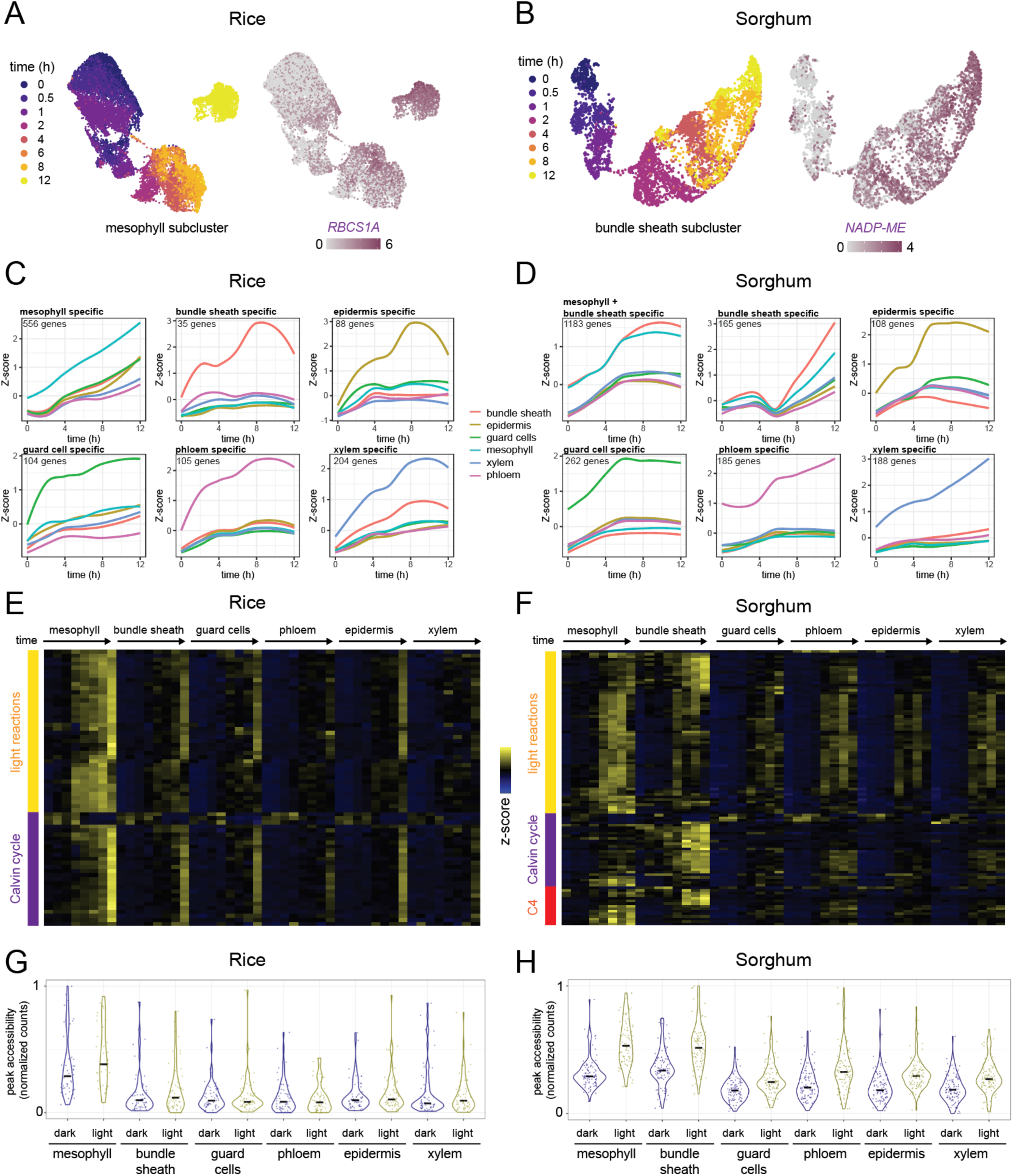
Light induces changes in cell-type specific transcript abundance and chromatin accessibility in rice and sorghum. **(A)** Sub-clustering of single nuclei transcript profiles from rice mesophyll during de-etiolation. Transcript abundance of *RbcS1A* gene shown on right. **(B)** Sub-clustering of single nuclei transcript profiles from sorghum bundle sheath during de-etiolation. Transcript abundance of *NADP-ME* shown on right. **(C & D)** Changes in transcript abundance from light responsive genes in the first 12h of exposure to light. Each cluster shows patterns of gene expression induction unique to each cell type. Clusters of genes were identified using Pearson correlation. **(E & F)** Heatmap derived from transcript abundance of photosynthesis genes in different cell types of rice and sorghum during the first 12h of exposure to light. **(G & H)** Normalized chromatin accessibility for photosynthesis genes in each cell type of rice and sorghum shoots at 0h (dark) and 12h (light).

A more detailed interrogation of the data showed an increase in transcript abundance of the light signaling transcription factor ELONGATED HYPOCOTYL 5 (HY5) (Lee et al., 2007), with a particularly strong response in guard cells of both rice and sorghum (**Figure S9**). Moreover, PHYTOCHROME INTERACTING FACTOR (PIF) transcription factors that repress photomorphogenesis in the dark (Moon et al., 2008; Gommers & Monte, 2018) were rapidly downregulated in response to light in all cell types (**Figure S9**). Light-induced partitioning of canonical photosynthesis genes between mesophyll and bundle sheath was also apparent (**Figure 5E, Table S8**). In rice, photosynthesis genes were most strongly induced in the mesophyll although a similar but weaker response was also seen in bundle sheath, guard, phloem, epidermis and xylem nuclei (**Figure 5E**). SEM confirmed that in the dark etioplasts were present in vascular and epidermal cells, and after exposure to light thylakoid-like membranes were evident (**Figure S10**). This supports the observation that photosynthesis can be weakly induced in these cell types. In sorghum, light strongly induced photosynthesis genes in both mesophyll and bundle sheath cell types, and included genes important for the light-dependent reactions of photosynthesis as well as the Calvin-Benson-Bassham and C_4_ cycles (**Figure 5F**). As expected, Calvin-Benson-Bassham cycle genes and *NADP-ME* were highly induced in sorghum bundle sheath, while *CARBONIC ANHYDRASE* and *PYRUVATE,ORTHOPHOSPHATE DIKINASE* were induced in mesophyll cells (**Table S8**). Agreeing with these data, chromatin of photosynthesis genes was more accessible in mesophyll cells compared with other cell types (t-test *p* < 5.9 x 10^-5^) (**Figure 5G**), however the difference in accessibility in response to light was only marginally significant in the rice mesophyll (*p* = 0.097). In contrast, in sorghum accessibility of photosynthesis genes increased in response to light in both mesophyll and bundle sheath cells (*p* < 2.9 x 10^-3^) (**Figure 5H**). These data indicate a pervasive gain of light regulation by photosynthesis genes in the bundle sheath of sorghum likely facilitated by increased chromatin accessibility.

### Cell identity conditions partitioning of photosynthesis gene expression in the dark

In both rice and sorghum differences in expression of photosynthesis genes between mesophyll and bundle sheath cells increased with time (**Figure 5E & 5F, Figure S11**). As would be expected, in rice these genes were preferentially expressed in mesophyll cells (**Figure 6B**) whilst in sorghum some were preferential to mesophyll cells and others were more highly expressed in the bundle sheath (**Figure 6B**). This is exemplified by the *GLYCOLATE OXIDASE* gene, whose transcripts showed greater partitioning to mesophyll cells of rice and bundle sheath cells of sorghum in response to light (**Figure 6A**). After 12h of light exposure 72 photosynthesis genes in rice and 77 in sorghum were partitioned between mesophyll and bundle sheath cells (**Figure 6B**). However, for some photosynthesis genes differences in expression between cells were evident in the dark. This suggests that cell identity conditions light responses. Specifically, in the dark 29% and 58% of photosynthesis transcripts in rice and sorghum respectively were significantly partitioned between mesophyll and bundle sheath cells (**Figure 6B**). For example, at 0 hours *Rbcs2* transcripts were already more abundant in mesophyll and bundle sheath cells of rice and sorghum respectively (**Figure 6A**). This finding is consistent with the fact that promoters of photosynthesis genes contained regions of open chromatin in the etiolated state (**Figure 5G, 5H**). Indeed, differences in accessible chromatin between mesophyll and bundle sheath cells at 0 hours were also evident in promoter regions of *GLYCOLATE OXIDASE* and *RBCS2* (**Figure 6C**). In fact, in the etiolated state many photosynthesis genes showed differences in chromatin accessibility between cell types (**Figure 6D**). In the dark open chromatin upstream of photosynthesis genes was predominantly found in mesophyll cells of rice and this was reinforced after exposure to light (**Figure 6D**). In sorghum open chromatin was evident upstream of canonical photosynthesis genes as well as those of the C_4_ cycle in both mesophyll and bundle sheath cells, and as in rice this was strengthened by light. We conclude that intrinsic differences in cell identity contribute to the partitioning of photosynthesis gene expression between cells in both C_3_ rice and C_4_ sorghum, and that differential partitioning is not driven exclusively by light signaling.

**Figure 6:**
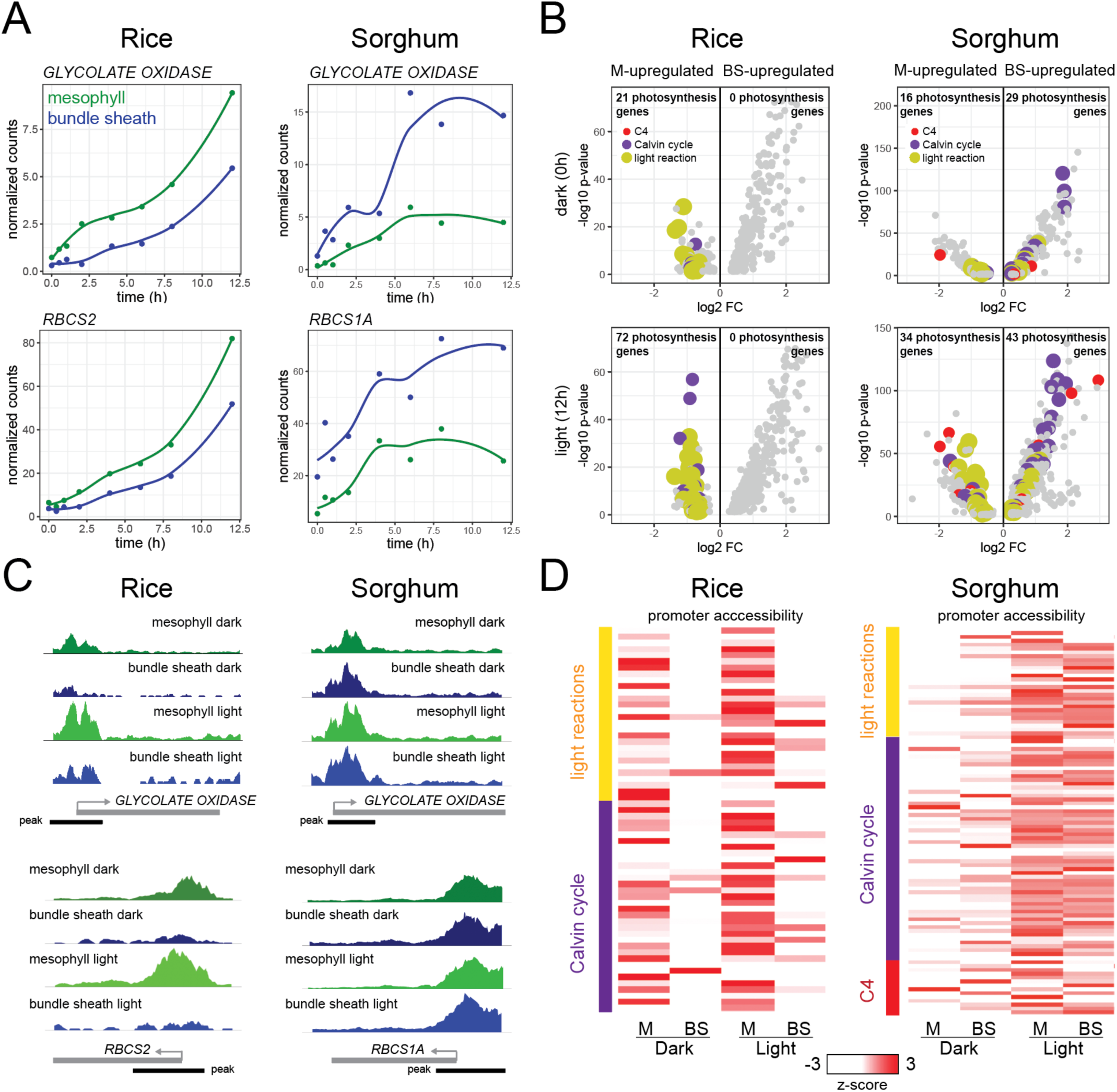
Cell identity drives partitioning of photosynthesis genes between mesophyll and bundle sheath cells in rice and sorghum. **(A)** Transcript abundance of *GLYCOLATE OXIDASE* and *RBCS2* during de-etiolation in mesophyll and bundle sheath cells of rice and sorghum. **(B)** Volcano plots of significantly differentially expressed genes between mesophyll and bundle sheath cells at 0h and 12h after exposure to light in rice and sorghum. Genes encoding enzymes involved in C_4_ photosynthesis, the Calvin Benson Bassham cycle and the light reactions are shown in red, purple, and yellow, respectively. **(C)** Chromatin accessibility in the mesophyll and bundle sheath promoters of *GLYCOLATE OXIDASE* and *RBCS2* subunit under etiolated and light conditions. **(D)** Chromatin accessibility differences adjacent (+-2000 bp) to photosynthesis genes at 0h (dark) and 12h (light) after light exposure.

### Photosynthesis genes in C_4_ sorghum acquire a *cis*-code associated with the C_3_ bundle sheath of rice

*Cis*-regulatory DNA sequences drive the patterning of gene expression (Marand et al., 2017; Kaufmann et al., 2010). Therefore, we next searched for *cis*-elements that underlie our observed cell identity - and light-dependent patterns of gene expression in rice and sorghum. When regions of open chromatin specific to each cell type were assessed for over-represented transcription factor binding sites, this identified the same motifs in the same cell types of both rice and sorghum (**Figure 7A, Figure S12, Table S9**). Thus, both species share a conserved cell-type specific *cis*-regulatory code. For example, motifs bound by Myeloblastosis (Myb)-related and NAM, ATAF1/2, and CUC2 (NAC) transcription factors defined accessible chromatin regions in mesophyll nuclei from both rice and sorghum, while the DOF motif was enriched in bundle sheath and phloem-specific peaks of both species (**Figure 7A, Figure S12**). Further, xylem-specific peaks were enriched in MYB and ANAC transcription factor motifs while peaks associated with epidermis and guard cells contained binding sites for the homeodomain GLABROUS 1 (HDG1), Zinc finger-homeodomain (ZHD) and AT-hook motif nuclear-localized (AHL) families of transcription factors. In contrast, when we examined motifs in chromatin that were differentially accessible in response to light, we found that the same motifs were enriched, regardless of cell type. These motifs comprised the light-responsive circadian clock related basic leucine zipper (bZIPs) and CIRCADIAN CLOCK ASSOCIATED 1 (CCA1) motifs (**Figure 7B, Figure S13, Table S10**). These findings suggest that cell-type specific patterning of gene expression is defined by cell-identity *cis*-elements, whereas light-responsive gene expression is regulated by similar *cis*-elements across all cell types.

**Figure 7:**
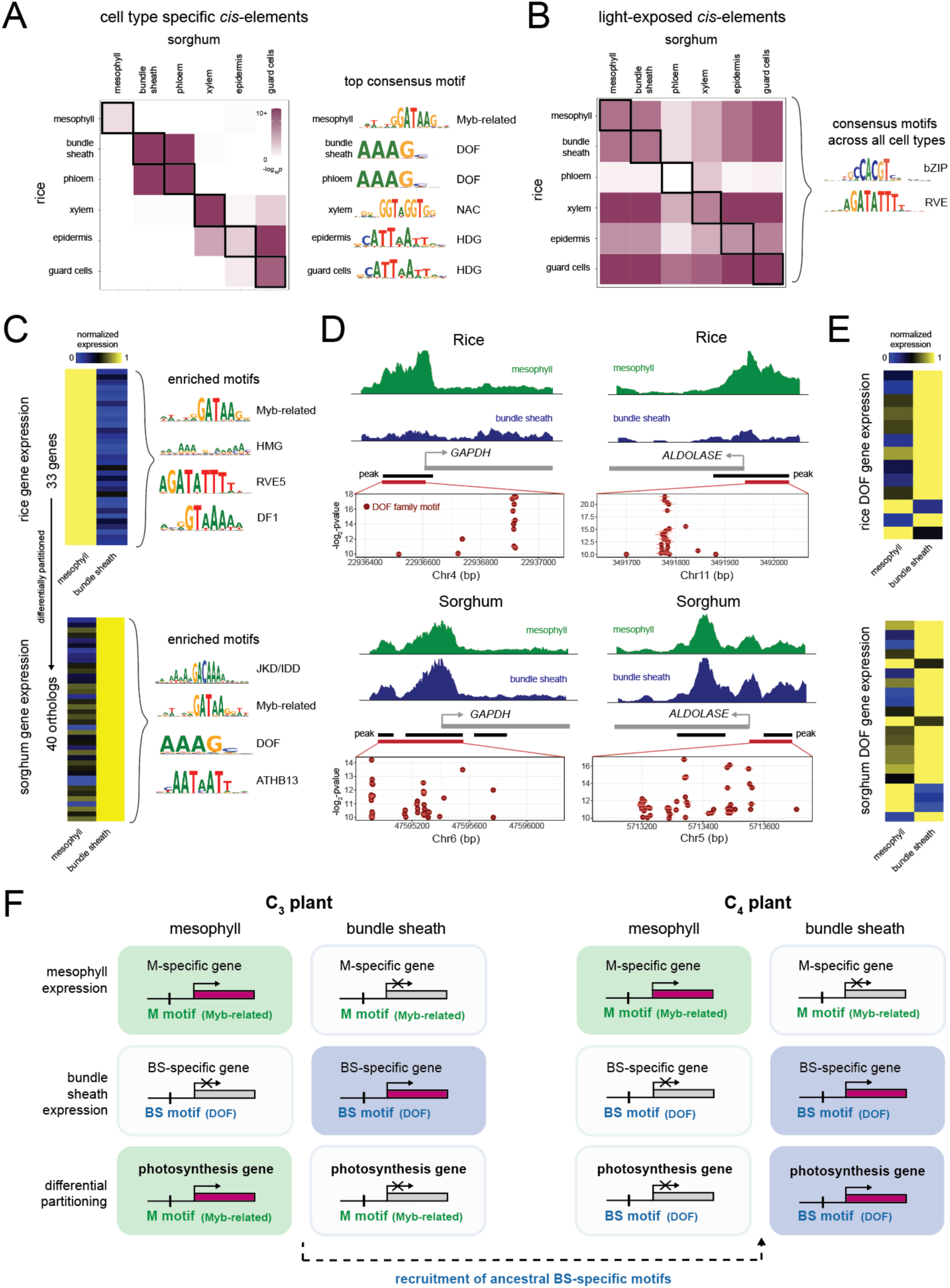
The cistrome of each cell type in C_3_ rice and C_4_ sorghum is conserved and drives partitioning of photosynthesis between mesophyll and bundle sheath cells. **(A)** Statistical overlap of *cis*-elements associated with accessible chromatin in each cell type of rice and sorghum shoots. Consensus motif for the most over-represented *cis*-element for each cell type shown on right. **(B)** Statistical overlap of *cis*-elements associated with accessible chromatin in each cell type in response to light in rice and sorghum shoots. The consensus motif for the most over-represented *cis*-element for all cell types is shown on the right. **(C)** Gene expression heatmaps (left) of differentially partitioned genes in rice and sorghum, and their most enriched *cis*-elements (right) in accessible chromatin. **(D)** Mapping accessible chromatin and quantifying of DOF family motifs for differentially partitioned *GAPDH* and *FRUCTOSE BISPHOSPHATE ALDOLASE* genes. **(E)** DOF transcription factor family expression patterns in mesophyll and bundle sheath cell types in each species. **(F)** By acquiring DOF *cis*-regulatory elements, C_4_ genes co-opt an ancestral bundle sheath cell identity network that is common between both species.

We next investigated whether the cell-type specific *cis*-code regulates genes that are differentially partitioned. To this end, we examined genes that were strongly expressed in the rice mesophyll, and whose orthologs were partitioned to the sorghum bundle sheath (**Figure 7C**). Among the 40 orthologs in this category were the Calvin-Benson-Bassham cycle genes *FRUCTOSE BISPHOSPHATE ALDOLASE* and *GLYCERALDEHYDE 3-PHOSPHATE DEHYDROGENASE* (*GAPDH)*, photorespiration genes such as *GLYCOLATE OXIDASE*, and light reaction genes including the *LHCII* subunit (**Figure 7C**). To find accessible chromatin associated with these peaks, we correlated changes in gene expression with changes in accessibility. Strikingly, among these differentially partitioned genes, we found that associated chromatin was enriched in cell-type specific Myb-related, Auxin Response Factor 4 (ARF4) and REVEILLE 5 (RVE5) binding sites in mesophyll-specific genes in rice (**Figure 7C**), but enriched in cell-type specific DOF and JACKDAW (JKD) / Indeterminate Domain (IDD) binding sites in bundle-sheath specific orthologs in sorghum (**Figure 7C, Figure S14, Table S11**). This indicates that these orthologs swapped their partitioning from mesophyll to bundle sheath through changing identity-associated *cis*-regulatory motifs. Specifically, our data suggest that DOF transcription factor motifs were acquired by genes that swap expression from the mesophyll of C_3_ rice to the bundle sheath of C_4_ sorghum.

To investigate further, we examined canonical photosynthesis genes that were differentially partitioned from rice mesophyll to the sorghum bundle sheath for the prevalence of DOF motifs. This showed that accessible chromatin upstream of *GAPDH* in sorghum was enriched in DOF motifs compared to rice (**Figure 7D**). A similar enrichment of DOF motifs was seen in accessible chromatin of the sorghum bundle sheath for *FRUCTOSE BISPHOSPHATE ALDOLASE* (**Figure 7D**) and for *NADP-ME* (**Figure S15**). Furthermore, DOF family transcription factors were typically more strongly expressed in bundle sheath cells of both rice and sorghum (**Figure 7E**) suggesting that the patterning of these transcription factors has not changed during the transition from C_3_ to C_4._ Curiously, we found that while differentially partitioned genes in rice displayed large differences in chromatin accessibility between mesophyll and bundle sheath cell types, these differences were much smaller in sorghum. This suggests that while chromatin accessibility increases in the bundle sheath, accessibility is not necessarily lost in the mesophyll (**Figure S14**).

From our analysis we propose a model that explains the rewiring of cell-type specific regulation of photosynthesis genes in C_4_ leaves (**Figure 7F**). The model suggests that (***i***) the same mesophyll and bundle sheath specific *cis*-elements are active in rice and sorghum; (***ii***) patterning of transcription factors between the two species is relatively stable; and (***iii***) photosynthesis genes expressed in the bundle sheath of sorghum have acquired DOF *cis*-regulatory elements associated with bundle sheath cells in rice.

## Discussion

The rice and sorghum single nuclei gene expression and chromatin accessibility atlases presented here provide novel insights into the molecular signatures associated with each major leaf cell type. These atlases reveal the cell type specific processes that enable photosynthesis to occur as leaves develop in the presence of light and provide insight into how gene expression patterns are rewired to perform C_4_ photosynthesis.

### Signatures of major cell types in cereal leaves

Until recently, it has not been technically feasible to compare the transcriptional identities of each cell type in C_3_ and C_4_ leaves. Our data provide a new level of granularity that allows us to explore the transcriptional identity of each cell type within a leaf and across two species. Our findings highlight the differences in functions that each cell type carries out. For example, specific gene expression in epidermal cells provides evidence for their role in the biosynthesis and export of cutin molecules used in the formation of the protective cuticle at the leaf surface, while cells associated with the phloem show high expression of genes involved in the transport of sugars and solutes. These data are consistent with previous single-cell RNA sequencing studies focusing on leaf tissues from *Arabidopsis thaliana* and rice (Procko et al., 2022; Berrío et al., 2022; Wang et al., 2021). By comparing transcriptional identities between cell types of rice and sorghum leaves, we found that the C_4_ sorghum bundle sheath has simultaneously gained novel expression of many genes, and as would be expected this included C_4_ genes. However, it was also apparent that the sorghum bundle sheath cells have lost expression of genes specific to the C_3_ bundle sheath. Notably, the sorghum bundle sheath acquired specific gene expression from other C_3_ cell types, with the C_3_ mesophyll and guard cells being prominent cell types that have swapped gene expression with the C_4_ bundle sheath. These findings have important implications for ongoing efforts to engineer C_3_ plants with C_4_ photosynthetic characteristics (Ermakova et al., 2021) and suggest that the C_3_ bundle sheath undergoes a broad change in transcriptional identity to achieve a functional C_4_ pathway. Our data also suggest that other cell types, such as the phloem and the guard cells, have been re-functionalized such that they carry out functions to support C_4_ photosynthesis in leaves.

### Establishing photosynthesis in specific cell types

In contrast to mammals, plant transcriptomes are highly dynamic in response to external environmental cues. To capture such changes, our gene expression atlases not only explore transcriptional identity, but also how cell transcriptional states change after the perception of light. Previous analysis of de-etiolation has proved a powerful system to understand the complex dynamics associated with photosynthesis (Boffey et al., 1980; Sullivan et al., 2014; Armarego-Marriott et al., 2019; Armarego-Marriott et al., 2020; Singh et al., 2023). Such studies have highlighted the extensive transcriptional and translational changes required for the assembly of chloroplast membranes and the photosynthetic apparatus, which in turn have elucidated the central role that light signaling plays in both chloroplast and seedling development. Moreover, several transcription factors, including positive regulators such as Hy5 and negative regulators such as PIFs, have been shown to play critical roles in regulating the expression of photosynthesis genes during de-etiolation (Moon et al., 2008; Shin et al., 2009; Leivar & Quail, 2011; Gommers & Monte, 2018). However, bulk analysis of leaf tissue has occluded the multitude of light-induced responses taking place in each cell type. By mapping gene expression at single cell resolution in rice and sorghum shoots at different time points during de-etiolation, we captured how the transcriptional state of each cell type responds to light and discovered both shared as well as unique gene expression responses. We found that all cell types activated photosynthesis gene expression to some extent and that light signaling-related *cis*-regulatory elements bZIP and CCA1 were differentially accessible in each cell type in response to light. However, there were clear differences in the magnitude and dynamics of photosynthesis gene induction. For example, vasculature cell types typically displayed weak activation, whereas the mesophyll in rice and both the mesophyll and bundle sheath in sorghum reacted strongly to light exposure. This indicates that light responsive gene expression is a generic feature of all cell types in a leaf, but it appears that the amplitude of the light response is conditioned by cell identity.

There is clear evidence that transcriptional reprogramming is critical after perception of light to induce chloroplast development and photosynthesis (Arsovski et al., 2012). Despite this, some studies show that cell identity can also affect photosynthesis gene expression (Langdale 1988; Wang et al., 1993). For example, transcripts of both the large and the small subunits of RuBisCO in the C_4_ plant *Amaranthus hypochondriacus* were found to be localized preferentially to the bundle sheath cells in cotyledons of dark-grown seedlings, while RuBisCO, PEPC and PPDK proteins were present in a cell type specific manner in the absence of light (Wang et al., 1993). Our observations support and expand these early reports to other members of the photosynthesis pathway. While light responses in mesophyll and bundle sheath cells in both rice and sorghum played an important role in the regulation of photosynthesis, our data show that partitioning of photosynthesis, both at the gene expression and chromatin level, was already present in the dark (i.e. in a light-independent manner). From these data, we conclude that light-independent, cell identity-driven partitioning of photosynthesis genes in mesophyll and bundle sheath cells occurs in both C_3_ and C_4_ plants and is a common phenomenon associated with many photosynthesis genes.

### C_4_ gene expression exapts gene regulatory networks associated with cell identity in C_3_ plants

Previous studies have identified a small number of *cis*- and *trans*-regulatory mechanisms that control the specific expression of C_4_ genes in mesophyll and bundle sheath cells (Marshall et al., 1997; Gowik et al., 2004; Akyildiz et l., 2007; Brown et al., 2011; Williams et al., 2016; Reyna-Llorens et al., 2018; Borba et al., 2023). The expression patterns driven by these *cis*-regulatory sequences in C_3_ and C_4_ species suggest that rewiring of ancestral gene regulatory mechanisms underlies the evolution of cell type specific gene expression required for C_4_ photosynthesis. However, until this point, efforts to understand cell type specific gene expression in leaves have primarily focused on C_4_ species, and so little is known about the ancestral *cis*-regulatory sequences and transcription factors that regulate gene expression in mesophyll or bundle sheath cells of genes from C_3_ species. To our knowledge a MYC and MYB bipartite transcription factor module is the only known gene regulatory mechanism that drives bundle sheath expression in a C_3_ plant (Dickinson et al., 2020). Here, a comparison between six major leaf cell types of rice and sorghum revealed that most cell types held distinct cell identity related *cis*-elements. Notably, there was a high degree of conservation of cell identity-associated motifs between rice and sorghum. For example, DOF transcription factor binding sites were enriched in chromatin regions that were specifically accessible in bundle sheath cells of both rice and sorghum. Supporting this, transient expression assays in C_4_ maize have shown that DOF transcription factors are able to activate the expression of *NADP-ME* and *PEPCK,* two genes that are specifically expressed in the bundle sheath (Borba et al., 2023). Thus, in contrast to the low level of conservation of transcriptional identities detected for the bundle sheath between C_3_ rice and C_4_ sorghum, it appears that the *cis*-regulatory landscape regulating cell-type specific expression is conserved between the two species. These results suggest that to change spatial patterns of gene expression within a leaf, genes recruit ancestral cell-type specific *cis*-regulatory mechanisms to re-direct gene expression to a new cell type. Our findings suggest that the bundle sheath specific regulation of C_4_ genes by DOF transcription factors might be a mechanism employed by several C_4_ species, and that this gene regulatory mechanism was recruited from the ancestral C_3_ state, a hypothesis that is supported by the observed enrichment of DOF motifs in bundle sheath specific accessible chromatin of C_3_ rice.

Various models have been proposed to explain the repeated evolution of the complex C_4_ phenotype. All are founded on analysis of genera such as *Flaveria* and *Cleome* that contain not just C_3_ and C_4_ species but also C_3_-C_4_ (or C_2_) species that show partial C_4_-like features such as reduced photorespiration and partial or complete Kranz anatomies (Marshall et al., 2007; Ku et al. 1991). A conceptual model based on observation orders events from the ancestral C_3_ to the derived C_4_ state (Sage et al., 2004; 2012). Here, C_4_ evolution has been proposed to have occurred in a stepwise manner involving changes in vein density, increased volume of the bundle sheath chloroplast compartment, loss of photorespiration in mesophyll cells and then activation of the full C_4_ cycle evolving consecutively (Sage et al., 2004; 2012). A statistical model using phenotypic landscape inference later indicated that the ordering of such events likely differs between lineages and thus that the C_4_ state is accessible from multiple routes derived from the C_3_ system (Williams et al., 2013). Lastly, a mathematical model predicts that once changes to C_3_ leaves take place such that C_2_ metabolism is initiated, a smooth Mount Fuji-like landscape leads to the full C_4_ pathway through gradual improvements in photosynthetic efficiency (Heckmann et al., 2013). In the future, these genera containing intermediate species will be ideal models to fully understand how C_4_ evolution has rewired gene patterning in all cell types of the leaf.

## Methods

### Plant growth

For the de-etiolation time course, seeds of *Oryza sativa spp. japonica* cultivar Kitaake and *Sorghum bicolor* BTx623 were incubated in sterile water for 2 days and 1 day respectively at 29°C in the dark. Germinated seedlings were transferred in a dark room equipped with green light to a 1:1 mixture of topsoil and sand supplemented with fertilizer granules and grown for 5 days in the dark by wrapping the tray and lid several times with aluminum foil. Plants were placed in a controlled environment room with 60% humidity, temperatures of 28°C and 20°C during the day and night, respectively. Plants were exposed to light at the beginning of a photoperiod of 12 h light and 12 h dark (Figure 1A), and shoots were harvested at different time points during de-etiolation by flash-freezing tissue in liquid nitrogen. For the 0 h time point, seedlings were harvested in a dark room equipped with green light and flash-frozen immediately.

For microscopy analysis and enrichment of bundle sheath nuclei using fluorescence-activated nuclei sorting, *Oryza sativa spp. japonica* cultivar Kitaake single copy homozygous T2 seeds were dehusked and sterilized in 10% (v/v) bleach for 30 min. After washing several times with sterile water, seeds were incubated for 2 days in sterile water at 29°C in the dark. Germinated seedlings were transferred to ½ strength Murashige and Skoog medium with 0.8% agar in magentas and grown for 5 days in the light in a growth chamber at temperatures of 28°C and 20°C during the day and night, respectively, and a photoperiod of 12 h light and 12 h dark.

### Rice transformation

*Oryza sativa* spp. *japonica* cultivar Kitaake was transformed using *Agrobacterium tumefaciens* as described previously (Hiei & Komari, 2008) with several modifications. Seeds were de-husked and sterilized with 10% (v/v) bleach for 15 min before placing them on nutrient broth (NB) callus induction medium containing 2 mg/L 2,4-dichlorophenoxyacetic acid for 4 weeks at 28°C in the dark. Growing calli were co-incubated with *A. tumefaciens* strain LBA4404 carrying the expression plasmid of interest in NB inoculation medium containing 40 μg/ml acetosyringone for 3 days at 22°C in the dark. Calli were transferred to NB recovery medium containing 300 mg/L timentin for 1 week at 28°C in the dark. They were then transferred to NB selection medium containing 35 mg/L hygromycin B for 4 weeks at 28°C in the dark. Proliferating calli were subsequently transferred to NB regeneration medium containing 100 mg/L myo-inositol, 2 mg/L kinetin, 0.2 mg/L 1-naphthaleneacetic acid, and 0.8 mg/L 6-benzylaminopurine for 4 weeks at 28°C in the light. Plantlets were transferred to NB rooting medium containing 0.1 mg/L 1-naphthaleneacetic acid and incubated in Magenta pots for 2 weeks at 28°C in the light. Finally, plants were transferred to a 1:1 mixture of topsoil and sand and grown in a controlled environment room with 60% humidity, temperatures of 28°C and 20°C during the day and night, respectively, and a photoperiod of 12-hr light and 12-hr dark.

### Construct design and cloning

The coding sequence for mTurquoise2 was obtained from Luginbuehl et al., 2020. The promoter sequence from *Zoysia japonica PHOSPHOENOLPYRUVATE CARBOXYKINASE* in combination with the dTALE STAP4 system was obtained from Danila et al., 2022. The coding sequence of *Arabidopsis thaliana* H2B (At5g22880) was used as an N-terminal signal for targeting mTurquoise2 to the nucleus. All sequences were domesticated for Golden Gate cloning (Engler et al., 2009; Weber et al., 2011). Level 1 and Level 2 constructs were assembled using the Golden Gate cloning strategy to create a binary vector for expression of *STAP4*-mTurquoise2-H2B driven by *PCK*-dTALE.

### Microscopy

To test bundle-sheath specific expression of mTurquoise2-H2B, recently expanded leaf 3 of 7-day old seedlings was prepared for confocal microscopy by scraping the adaxial side of the leaf blade two to three times with a sharp razor blade, transferring to water to avoid drying out and then mounting on a microscope slide with the scraped surface facing upwards. Confocal imaging was performed on a Leica TCS SP8 X using a 10X air objective (HC PL APO CS2 10X 0.4 Dry) with optical zoom, and hybrid detectors for fluorescent protein and chlorophyll autofluorescence detection. The following excitation (Ex) and emission (Em) wavelengths were used for imaging: mTurquoise2 (Ex = 442, Em = 471–481), chlorophyll autofluorescence (Ex = 488, Em = 672–692).

### Enrichment of bundle sheath nuclei using fluorescence-activated cell sorting

To purify the nuclei population from whole leaves, recently expanded leaves 3 from five 7-day old wild type rice seedlings were chopped on ice in Nuclei Buffer (10 mM Tris-HCl, pH 7.4, 10 mM NaCl, 3 mM MgCl_2_, 0.5 mM spermidine, 0.2 mM spermine, 0.01% Triton-X, 1x Roche complete protease inhibitors, 1% BSA, Protector RNase inhibitor) with a sharp razor blade. The suspension was filtered through a 70 um filter and subsequently through a 35 μm filter. Nuclei were stained with Hoechst and FACS purified on an AriaIII instrument, using a 70 μm nozzle. Nuclei were collected in an Eppendorf tube containing BSA and Protector RNase inhibitor. Using the same approach, nuclei from the bundle sheath marker line expressing mTurquoise2-H2B were isolated. Nuclei were sorted based on the mTurquoise2 fluorescent signal. Nuclei were collected in minimal Nuclei Buffer (10 mM Tris-HCl, pH 7.4, 10 mM NaCl, 3 mM MgCl_2_, RNase inhibitor, 0.05% BSA). After collection, nuclei were spun down in a swinging bucket centrifuge at 405g for 5 mins, with reduced acceleration and deceleration. Nuclei were resuspended in minimal Nuclei Buffer and mixed with the unspun whole leaf nuclei population to achieve a proportion of approximately 25% mTurquoise2-positive nuclei. The bundle sheath enriched nuclei population was sequenced using the 10X Genomics Gene Expression platform with v3.1 chemistry and sequenced on the Illumina Novaseq 6000 with 150 bp-paired end chemistry.

### Chlorophyll quantification

Seedlings were harvested at specified time points during de-etiolation and immediately flash-frozen in liquid nitrogen. Frozen tissue was ground into fine powder and the weight was measured before suspending the tissue in 1 ml of 80% (v/v) acetone. After vortexing, the tissue was incubated on ice for 15 min with occasional mixing of the suspension. The tissue was spun down at 15,700 g at 4°C and the supernatant was removed. The extraction was repeated and supernatants were pooled before measuring the absorbance at 663.6nm and 646.6nm in a spectrophotometer. The total chlorophyll content was determined as described previously (Porra et al., 1989).

### Scanning electron microscopy

For the de-etiolation experiment of rice and sorghum, samples from 4-6 individual seedlings for each time point (0 h, 6 h, 12 h, 48 h) were harvested for electron microscopy. Leaf segments (∼2 mm^2^) were excised with a razor blade and immediately fixed in 2% (v/v) glutaraldehyde and 2% (w/v) formaldehyde in 0.05 - 0.1 M sodium cacodylate (NaCac) buffer (pH 7.4) containing 2 mM calcium chloride. Samples were vacuum infiltrated overnight, washed 5 times in 0.05 – 0.1 M NaCac buffer, and post-fixed in 1% (v/v) aqueous osmium tetroxide, 1.5% (w/v) potassium ferricyanide in 0.05 M NaCac buffer for 3 days at 4°C. After osmication, samples were washed 5 times in deionized water and post-fixed in 0.1% (w/v) thiocarbohydrazide for 20 min at room temperature in the dark. Samples were then washed 5 times in deionized water and osmicated for a second time for 1 h in 2% (v/v) aqueous osmium tetroxide at room temperature. Samples were washed 5 times in deionized water and subsequently stained in 2% (w/v) uranyl acetate in 0.05 M maleate buffer (pH 5.5) for 3 days at 4°C and washed 5 times afterwards in deionized water. Samples were then dehydrated in an ethanol series, transferred to acetone, and then to acetonitrile. Leaf samples were embedded in Quetol 651 resin mix (TAAB Laboratories Equipment Ltd) and cured at 60°C for 2 days. Ultra-thin sections of embedded leaf samples were prepared and placed on Melinex (TAAB Laboratories Equipment Ltd) plastic coverslips mounted on aluminum SEM stubs using conductive carbon tabs (TAAB Laboratories Equipment Ltd), sputter-coated with a thin layer of carbon (∼30 nm) to avoid charging and imaged in a Verios 460 scanning electron microscope at 4 keV accelerating voltage and 0.2 nA probe current using the concentric backscatter detector in field-free (low magnification) or immersion (high magnification) mode (working distance 3.5 – 4 mm, dwell time 3 µs, 1536 x 1024 pixel resolution). For overserving plastid ultrastructure, SEM stitched maps were acquired at 10,000X magnification using the FEI MAPS automated acquisition software. Greyscale contrast of the images was inverted to allow easier visualisation.

### Nuclei extraction and single nuclei RNA sequencing (10X RNA-seq)

Frozen tissue from each time point (1 biological replicate per time point, 8 time points) was crushed using a bead bashing approach, and nuclei released from homogenate by resuspending in Nuclei Buffer (10 mM Tris-HCl, pH 7.4, 10 mM NaCl, 3 mM MgCl2). The resulting suspension was passed through a 30-um filter. To enrich the filtered solution for nuclei, an Optiprep (Sigma) gradient was used. Enriched nuclei were then stained with Hoechst, before being FACS purified. Purified nuclei were run on the 10X-Gene Expression platform with v3.0 chemistry, and sequenced on the Illumina Novaseq 6000 with 150 bp paired end chemistry. Single cell libraries were made following the manufacturers protocol. Libraries were sequenced to an average saturation of 63 % (14 % s.d.) and aligned either to the rice (Oryza sativa, subspecies *Nipponbare*; MSU annotation) (Kawahara et al., 2013) or sorghum genome (Sorghum bicolor v3.0.1; JGI annotation; McCormick et al., 2018). Chloroplast and mitochondrial reads were removed. For each time point, an average of 12,524 nuclei were sequenced (6,405 s.d.), with an average median UMI of 1,152 (420 s.d.) across both species. Doublets were removed using doubletFinder (McGinnis et al., 2019).

### Nuclei extraction and single nuclei RNA sequencing (sciRNA-seq3)

Each individual frozen seedling (10 - 12 individual seedlings per time point) was crushed using a bead bashing approach in 96 well plate, after which homogenate was resuspended in Nuclei Buffer. Resulting suspensions were passed through a 30-um filter. Washed nuclei were then reverse-transcribed with a well specific primer. After this step, remaining pool and split steps for sci-RNA-seq3 were followed as outlined in (Cao et al., 2019). We note the same approach was used to sequence the 48 hr time point, however a population of 6 plants were used instead of individual seedlings. Libraries were sequenced to an average saturation of 80 % (5 % s.d.), and sequenced on the Illumina Novaseq 6000 with 150 bp paired-end chemistry. Reads and aligned to either the rice or sorghum genome as described above. Chloroplast and mitochondrial reads were removed. For 0 - 12 h timepoints, an average of 6,527 nuclei were sequenced (5,039 s.d.), with an average median UMI of 423 (41 s.d.) across both species. For the 48 h time point, 77,208 and 82,748 nuclei were sequenced with a median UMI of 757 and 740 for rice and sorghum respectively.

### Nuclei extraction and single nuclei RNA sequencing (10X Multiome)

Fresh seedling tissue was harvested after 0 or 12 h light treatment (2 biological replicates per species, each with 2 - 4 technical replicates per time point, n = 11). Fresh tissue was chopped finely on ice in green room conditions in Nuclei Buffer. The resulting homogenate was filtered using a 30-um filter. Nuclei were enriched using Optiprep gradient. No FACS was performed. Nuclei were run on the 10X-Multiome platform with v1.0 chemistry. Single cell libraries made following manufacturers protocol, and sequenced on the Illumina Novaseq 6000with 150 bp paired-end chemistry. Reads and aligned to either the rice or sorghum genome as described above. Chloroplast and mitochondrial reads were removed. For each sample, an average of 1,923 nuclei were sequenced (1,334 s.d.), with an average median UMI of 1,644 (646 s.d.) and median ATAC fragments 10,251 (7,001 s.d.) across both species.

### Nuclei Clustering

Transcriptional atlases were generated separately for each species using Seurat (Hao, Hao et al., 2021). Nuclei were aggregated across various time points (ranging from 0 to 48 h) and methods (10X and sci-RNA-seq3). The integrated dataset was subjected to clustering, employing the top 2000 variable features that were shared across all datasets. Subsequent UMAP projections were constructed using the first 30 principal components. To analyse the rice bundle sheath specific mTurquoise line, we integrated two treatment replicates into a unified dataset. For this dataset, we clustered utilising the first 30 principal components. Cluster-specific markers were identified using the FindMarkers() command (adjusted p-value < 0.01). To determine the correspondence between the mTurquoise-positive cluster and clusters within the rice-RNA atlas, we compared the lists of cluster-specific markers (adjusted p-value < 0.01, specificity > 2) to those obtained from the rice-RNA atlas. For the 10X-multiome (RNA+ATAC) clustering we employed Signac (Stuart et al., 2021). Biological and technical replicates for each species were integrated, and clustering was conducted using the first 50 principal components derived from expression data. Following the initial peak calling using cellranger (10X genomics), peaks were subsequently re-called using MACS2 (Zhang et al., 2008). Differentially accessible peaks between cell types were identified using the FindMarkers() command (adjusted p-value < 0.05, percent threshold > 0.3).

### Orthology Analyses

We determined gene orthologs between rice and sorghum using OrthoFinder (Emms & Kelly, 2019). We constructed pan-transcriptome atlases by selecting expressed rice and sorghum genes that had cross-species orthologs. Ortholog conversions were performed in a one-to-one manner, meaning that if multiple orthologs for a gene were found across species, only one was retained. We integrated these datasets with Seurat using clustering approaches described above. To assign cell identities, we drew upon cell type labels that were previously assigned to each species separately and mapped them onto the pan-transcriptome clusters. To assess specific transcriptional differences in gene expression between the bundle sheath clusters of sorghum and rice within this dataset, we used the FindMarkers() command (adjusted p-value < 0.05). To examine the overlap of cell type-specific gene expression markers between the two species, we identified cell type markers from our main transcriptional dataset using FindMarkers() (adjusted p-value < 0.05, min.pct > 0.1). We note that some genes were found to be significant across multiple cell types. To assess the significance of this overlap, we compared their orthogroups and conducted a Fisher Exact Test, with the total number of orthogroups in the dataset as the background. Next, we assessed consistent and differential partitioning of gene expression patterns among each cell type pair (15 pairs total). To do this, we first calculated differentially expressed genes for each cell type pair by pseudo bulking transcriptomes of individual cell types across 0 - 12 h time points. Next, we identified partitioned expression patterns between cell types using an ANCOVA model implemented in DESeq2 (adj. *p* < 0.05). To perform cross-species comparisons of cell type pairs, we first converted differentially expressed genes to their Orthogroup. We then overlapped each cell type pair across species and evaluated the significance of these overlaps using the Fisher Exact Test. Finally, to distinguish whether a gene displayed consistent or differential partitioning in a particular cell type, we examined whether its fold change expression was higher or lower compared to its counterpart in the corresponding cell type of the other species.

### Differential Expression and Accessibility Responses to Light

We discovered cell-type specific differentially expressed genes during the first 12 h of light by pseudo-bulking transcriptional profiles. For each cell type, we then calculated the first and second principal component of these bulked profiles and found differentially expressed genes through fitting linear models to each of these principal components, as well as those that responded linearly with time using DESeq2 (adj. *p* < 0.05). To this list of differentially expressed genes, we also included genes that were differentially expressed between time point 0 and 12 in a pairwise test (adj. *p* < 0.05). Next, to uncover the different trends of gene expression among differentially expressed genes, we clustered genes using hierarchical clustering; choosing clustering cut offs that resulted in 10 rice and 18 sorghum clusters that contained at least 10 genes. To visualize the expression of these clusters, we scaled the expression of these clusters and fit a non-linear model to capture the dominant expression trend. Accessible chromatin within canonical photosynthesis genes were found through pseudo-bulking accessible chromatin by cell type. Accessible peaks needed to be within 2000 bp of the gene body and only 1 peak per gene is displayed. To compare peak accessibility across species, reads per peak were normalized between 0 and 1. Significant differences in accessibility between cell types of this group of genes were assessed using a paired t-test.

### *cis*-Element Analyses

To analyze over-represented cis-elements, we first calculated position frequency matrices using the JASPER2020 plant taxon group (Castro-Mondragon et al., 2021) with BSGenome assembled genomes (Pagès 2023). We found cell type specific accessible motifs per clustering using the RunchromVar function in Signac. This same approach was used for light responsive *cis*-elements, using light and dark treated nuclei within each cell type. We overlapped these cis-regulatory element lists by first sub-setting by the top 25 most significantly overrepresented motifs (adj. *p*-value < 0.05), before computing a Fisher Exact Test using all computed motifs as background. We clustered motifs using Tobias (Bentsen et al., 2020). To find differentially partitioned orthologous genes within our multiome gene expression dataset, we found mesophyll and bundle sheath specific genes in rice and sorghum respectively using the FindMarkers() command, with a p-value threshold cut off of 0.01 and a specificity above 1.25. To find overrepresented motifs within these genes, we correlated peak accessibility with gene expression using the LinkPeaks() command and kept only those peaks which were significantly associated with gene expression. We identified enriched cis-elements within these peaks using the FindMotifs() command; ranking by significance (adj. *p*-value < 0.05). We iterated the FindMotifs() command over 100 permutations to rank motifs that were consistently reported as enriched. Finally, to find DOF binding sites within accessible chromatin, we took the sequence underneath peaks upstream of target genes; ignoring areas that were inside gene bodies. We then used the runFimo() command with each DOF motifs that were found over-represented and conserved in the bundle sheath across species (adj. *p*-value < 0.01).

## Supplemental Information

Supplementary Figures (S1 – S15)

Supplementary Tables (S1 – S11)

Sequencing data can be found at the National Center for Biotechnology Information Sequence Read Archive, with Accession number PRJNAXXXXX.

## Acknowledgements

J.S. is an awardee of the Life Sciences Research Foundation (funded by Open Philanthropy) and a recipient of the American-Australian Association Fellowship (funded by Pratt Industries). L.H.L is a recipient of the Herchel Smith Postdoctoral Research Fellowship and was supported by BBSRC grant BBP0031171 to J.M.H. T.B.S was supported by a Swiss National Science Foundation (SNSF) Postdoc Mobility Fellowship (P500PB_203128) and an EMBO Long-Term Fellowship (ALTF 531-2019). J.R.E is an Investigator of the Howard Hughes Medical Institute. For the purpose of open access, the authors have applied a Creative Commons Attribution (CC BY) license to any Author Accepted Manuscript version arising from this submission. We thank Karin H. Müller, Georgina E. Lindop and Melissa J. Drignon from the Cambridge Advanced Imaging Centre for the electron microscopy sample preparation as well as support during the image acquisition.

## Author Contributions

L.H.L, J.S, J.R.E and J.M.H designed the experimental plan. J.S. and L.H.L performed laboratory experiments and genomic analyses. T.B.S. performed SEM imaging. R.M.D. carried out stable rice transformation. T.A.L optimized nuclei isolation. L.H.L., J.S., J.M.H., and J.R.E. wrote the manuscript, with input from all authors.

## References

Armarego-Marriott, T. et al. (2019). Highly resolved systems biology to dissect the etioplast-to-chloroplast transition in tobacco leaves. Plant Physiol. 180: 654–681.

Armarego-Marriott, T., Sandoval-Ibañez, O., and Kowalewska, Ł. (2020). Beyond the darkness: Recent lessons from etiolation and de-etiolation studies. J. Exp. Bot. 71: 1215–1225.

Aubry, S., Kelly, S., Kümpers, B.M.C., Smith-Unna, R.D., and Hibberd, J.M. (2014). Deep Evolutionary Comparison of Gene Expression Identifies Parallel Recruitment of Trans-Factors in Two Independent Origins of C4 Photosynthesis. PLoS Genet. 10: e1004365

Bassham, J.A., Benson, A.A., and Calvin, M. (1950). The path of carbon in photosynthesis. J. Biol. Chem. 185: 781–787.

Bauwe, H., Hagemann, M., and Fernie, A.R. (2010). Photorespiration: players, partners and origin. Trends Plant Sci. 15: 330–336.

Bentsen, M., Goymann, P., Schultheis, H., Klee, K., Petrova, A., Wiegandt, R., Fust, A., Preussner, J., Kuenne, C., Braun, T., Kim, J., and Looso, M. (2020). ATAC-seq footprinting unravels kinetics of transcription factor binding during zygotic genome activation. Nat. Commun. 11: 4267

Berrío, R.T., Verstaen, K., Vandamme, N., Pevernagie, J., Achon, I., van Duyse, J., van Isterdael, G., Saeys, Y., de Veylder, L., Inzé, D., and Dubois, M. (2022). Single-cell transcriptomics sheds light on the identity and metabolism of developing leaf cells. Plant Physiol. 188: 898–918.

Boffey, S.A., Sellden, G., and Leech, R.M. (1980). Influence of Cell Age on Chlorophyll Formation in Light-grown and Etiolated Wheat Seedlings. Plant Physiol. 65: 680–684.

Borba, A.R., Reyna-Llorens, I., Dickinson, P.J., Steed, G., Gouveia, P., Górska, A.M., Gomes, C., Kromdijk, J., Webb, A.A.R., Saibo, N.J.M., and Hibberd, J.M. (2023). Compartmentation of photosynthesis gene expression in C4 maize depends on time of day. Plant Physiol. 00: 1–15.

Bowes, G., Ogren, W.L., and Hageman, R.H. (1971). Phosphoglycolate production by Ribulose Diphosphate Carboxylase. Biochem. Biophys. Res. Commun. 45: 716–722.

Brown, N.J., Newell, C.A., Stanley, S., Chen, J.E., Perrin, A.J., Kajala, K., and Hibberd, J.M. (2011). Independent and Parallel Recruitment of Preexisting Mechanisms Underlying C4 Photosynthesis. Science 331: 1436–1439.

Burgess, S.J., Reyna-Llorens, I., Stevenson, S.R., Singh, P., Jaeger, K., and Hibberd, J.M. (2019). Genome-wide transcription factor binding in leaves from C3 and C4 grasses. Plant Cell 31: 2297– 2314.

Cao, J., Spielmann, M., Qiu, X., Huang, X., Ibrahim, D.M., Hill, A.J., Zhang, F., Mundlos, S., Christiansen, L., Steemers, F.J., Trapnell, C., and Shendure, J. (2019). The single-cell transcriptional landscape of mammalian organogenesis. Nature 566: 496–502.

Castro-Mondragon, J.A. et al. (2022). JASPAR 2022: The 9th release of the open-access database of transcription factor binding profiles. Nucleic Acids Res. 50: D165–D173.

Cervantes-Pérez, S.A., Thibivillliers, S., Tennant, S., and Libault, M. (2022). Review: Challenges and perspectives in applying single nuclei RNA-seq technology in plant biology. Plant Sci. 325: 1– 30.

Christin, P.A., Arakaki, M., Osborne, C.P., and Edwards, E.J. (2015). Genetic enablers underlying the clustered evolutionary origins of C4 photosynthesis in angiosperms. Mol. Biol. Evol. 32: 846– 858.

Covshoff, S., Furbank, R.T., Leegood, R.C., and Hibberd, J.M. (2013). Leaf rolling allows quantification of mRNA abundance in mesophyll cells of sorghum. J. Exp. Bot. 64: 807–813.

Danila, F. et al. (2022). A single promoter-TALE system for tissue-specific and tuneable expression of multiple genes in rice. Plant Biotechnol. J. 20: 1786–1806.

Emms, D.M. and Kelly, S. (2019). OrthoFinder: Phylogenetic orthology inference for comparative genomics. Genome Biol. 20: 1–14.

Engler, C., Gruetzner, R., Kandzia, R., and Marillonnet, S. (2009). Golden gate shuffling: a one-pot DNA shuffling method based on type IIs restriction enzymes. PLoS One 4: e5553.

Ermakova, M. et al. (2021). Installation of C4 photosynthetic pathway enzymes in rice using a single construct. Plant Biotechnol. J. 19: 575–588.

Furbank, R.T. (2011). Evolution of the C 4 photosynthetic mechanism: Are there really three C 4 acid decarboxylation types? J. Exp. Bot. 62: 3103–3108.

Ghannoum, O., Evans, J.R., and von Caemmerer, S. (2011). Nitrogen and water use efficiency of C4 plants. C4 Photosynth. Relat. CO2 Conc. Mech.: 129–146.

Gommers, C.M.M. and Monte, E. (2018). Seedling establishment: A dimmer switch-regulated process between dark and light signaling. Plant Physiol. 176: 1061–1074.

Gowik, U., Burscheidt, J., Akyildiz, M., Schlue, U., Koczor, M., Streubel, M., and Westhoff, P. (2004). cis-Regulatory Elements for Mesophyll-Specific Gene Expression in the C4 Plant Flaveria trinervia, the Promoter of the C4 PEPC Gene. Plant Cell 16: 1077–1090.

Grindberg, R. V. et al. (2013). RNA-sequencing from single nuclei. PNAS 110: 19802–19807.

Grosberg, R.K. and Strathmann, R.R. (2007). The evolution of multicellularity: A minor major transition? Annu. Rev. Ecol. Evol. Syst. 38: 621–654.

Guillotin, B., Rahni, R., Passalacqua, M., Mohammed, M.A., Xu, X., Raju, S.K., Ramírez, C.O., Jackson, D., Groen, S.C., Gillis, J., and Birnbaum, K.D. (2023). A pan-grass transcriptome reveals patterns of cellular divergence in crops. Nature 617: 785–791.

Haberlandt, G. (1884). Physiologische Pflanzenanatomie. W. Engelman.

Hao, Y. et al. (2021). Integrated analysis of multimodal single-cell data. Cell 184: 3573–3587.e29.

Hatch, M.D. and Slack, C.R. (1966). Photosynthesis by sugar-cane leaves. A new carboxylation reaction and the pathway of sugar formation. Biochem. J. 101: 103–11.

Heckmann, D., Schulze, S., Denton, A., Gowik, U., Westhoff, P., Weber, A.P.M., and Lercher, M.J. (2013). Predicting C4 photosynthesis evolution: Modular, individually adaptive steps on a mount fuji fitness landscape. Cell 153: 1579.

Hibberd, J.M. and Covshoff, S. (2010). The Regulation of Gene Expression Required for C _4_ Photosynthesis. Annu. Rev. Plant Biol. 61: 181–207.

Hiei, Y. and Komari, T. (2008). Agrobacterium-mediated transformation of rice using immature embryos or calli induced from mature seed. Nat. Protoc. 3: 824–834.

Hua, L., Stevenson, S.R., Reyna-Llorens, I., Xiong, H., Kopriva, S., and Hibberd, J.M. (2021). The bundle sheath of rice is conditioned to play an active role in water transport as well as sulfur assimilation and jasmonic acid synthesis. Plant J. 107: 268–286.

Huang, W. et al. (2022). A well-supported nuclear phylogeny of Poaceae and implications for the evolution of C4 photosynthesis. Mol. Plant 15: 755–777.

Jiao, Y., Ma, L., Strickland, E., and Deng, X.W. (2005). Conservation and divergence of light-regulated genome expression patterns during seedling development in rice and Arabidopsis. Plant Cell 17: 3239–3256.

John, C.R., Smith-Unna, R.D., Woodfield, H., Covshoff, S., and Hibberd, J.M. (2014). Evolutionary convergence of cell-specific gene expression in independent lineages of C4 grasses. Plant Physiol. 165: 62–75.

Jordan, D.B. and Ogren, W.L. (1984). The CO2/O2 specificity of ribulose 1,5-bisphosphate carboxylase/oxygenase : Dependence on ribulosebisphosphate concentration, pH and temperature. Planta 161: 308–313.

Kaufmann, K., Pajoro, A., and Angenent, G.C. (2010). Regulation of transcription in plants: Mechanisms controlling developmental switches. Nat. Rev. Genet. 11: 830–842.

Kawahara, Y. et al. (2013). Improvement of the oryza sativa nipponbare reference genome using next generation sequence and optical map data. Rice 6: 3–10.

Ku, M.S.B., Wu, J., Dai, Z., Scott, R.A., Chu, C., and Edwards, G.E. (1991). Photosynthetic and photorespiratory characteristics of Flaveria species. Plant Physiol. 96: 518–528.

Langdale, J.A., Zelitch, I., Miller, E., and Nelson, T. (1988). Cell position and light influence C4 versus C3 patterns of photosynthetic gene expression in maize. EMBO J. 7: 3643–3651.

Langdale, J.A. (2011). C4 Cycles: Past, present, and future research on C4 photosynthesis. Plant Cell 23: 3879–3892.

Lee, J., He, K., Stolc, V., Lee, H., Figueroa, P., Gao, Y., Tongprasit, W., Zhao, H., Lee, I., and Xing, W.D. (2007). Analysis of transcription factor HY5 genomic binding sites revealed its hierarchical role in light regulation of development. Plant Cell 19: 731–749.

Lee, T.A., Nobori, T., Illouz-Eliaz, N., Xu, J., Jow, B., Nery, J.R., and Ecker, J.R. (2023). A Single-Nucleus Atlas of Seed-to-Seed Development in Arabidopsis. bioRxiv: 2023.03.23.533992.

Luginbuehl, L.H., El-Sharnouby, S., Wang, N., and Hibberd, J.M. (2020). Fluorescent reporters for functional analysis in rice leaves. Plant Direct 4: 1–10.

Marand, A.P., Zhang, T., Zhu, B., and Jiang, J. (2017). Towards genome-wide prediction and characterization of enhancers in plants. Biochim. Biophys. Acta - Gene Regul. Mech. 1860: 131– 139.

Markelz, N.H., Costich, D.E., and Brutnel, T.P. (2003). Photomorphogenic Responses in Maize Seedling Development. Plant Physiol. 133: 1578–1591.

Marshall, D.M., Muhaidat, R., Brown, N.J., Liu, Z., Stanley, S., Griffiths, H., Sage, R.F., and Hibberd, J.M. (2007). Cleome, a genus closely related to Arabidopsis, contains species spanning a developmental progression from C3 to C4 photosynthesis. Plant J. 51: 886–896.

McCormick, R.F. et al. (2018). The Sorghum bicolor reference genome: improved assembly, gene annotations, a transcriptome atlas, and signatures of genome organization. Plant J. 93: 338–354.

McGinnis, C.S., Murrow, L.M., and Gartner, Z.J. (2019). DoubletFinder: Doublet Detection in Single-Cell RNA Sequencing Data Using Artificial Nearest Neighbors. Cell Syst. 8: 329–337.e4.

Moon, J., Zhu, L., Shen, H., and Huq, E. (2008). PIF1 directly and indirectly regulates chlorophyll biosynthesis to optimize the greening process in Arabidopsis. PNAS 105: 9433–9438.

Nobori, T., Monell, A., Lee, T.A., Zhou, J., Nery, J., and Ecker, J.R. (2023). Time-resolved single-cell and spatial gene regulatory atlas of plants under pathogen attack. bioRxiv: 2023.04.10.536170.

Pagès H. (2023). BSgenome: Software infrastructure for efficient representation of full genomes and their SNPs. R package version 1.68.0, https://bioconductor.org/packages/BSgenome.

Porra, R.J., Thompson, W.A., and Kriedemann, P.E. (1989). Determination of accurate extinction coefficients and simultaneous equations for assaying chlorophylls a and b extracted with four different solvents: verification of the concentration of chlorophyll standards by atomic absorption spectroscopy. Biochim. Biophys. Acta: 384–394.

Procko, C., Lee, T., Borsuk, A., Bargmann, B.O.R., Dabi, T., Nery, J.R., Estelle, M., Baird, L., O’Connor, C., Brodersen, C., Ecker, J.R., and Chory, J. (2022). Leaf cell-specific and single-cell transcriptional profiling reveals a role for the palisade layer in UV light protection. Plant Cell 34: 3261–3279.

Reyna-Llorens, I., Burgess, S.J., Reeves, G., Singh, P., Stevenson, S.R., Williams, B.P., Stanley, S., and Hibberd, J.M. (2018). Ancient duons may underpin spatial patterning of gene expression in C 4 leaves. PNAS 115: 1931–1936.

Sage, R.F. (2004). The evolution of C 4 photosynthesis. New Phytol. 161: 341–370.

Sage, R.F., Sage, T.L., and Kocacinar, F. (2012). Photorespiration and the evolution of C4 photosynthesis. Annu. Rev. Plant Biol. 63: 19–47.

Sage, R.F. and Zhu, X.G. (2011). Exploiting the engine of C 4 photosynthesis. J. Exp. Bot. 62: 2989– 3000.

Sheen, J.Y. and Bogorad, L. (1987). Differential expression of C4 pathway genes in mesophyll and bundle sheath cells of greening maize leaves. J. Biol. Chem. 262: 11726–11730.

Shin, J., Kim, K., Kang, H., Zulfugarov, I.S., Bae, G., Lee, C.H., Lee, D., and Choi, G. (2009). Phytochromes promote seedling light responses by inhibiting four negatively-acting phytochrome-interacting factors. PNAS 106: 7660–7665.

Singh, P., Stevenson, S.R., Dickinson, P.J., Reyna-Llorens, I., Tripathi, A., Reeves, G., Schreier, T.B., and Hibberd, J.M. (2023). C4 gene induction during de-etiolation evolved through changes in cis to allow integration with ancestral C3 gene regulatory networks. Sci. Adv. 9.

Stuart, T., Srivastava, A., Madad, S., Lareau, C.A., and Satija, R. (2021). Single-cell chromatin state analysis with Signac. Nat. Methods 18: 1333–1341.

Sullivan, A.M. et al. (2014). Mapping and dynamics of regulatory DNA and transcription factor networks in A. thaliana. Cell Rep. 8: 2015–2030.

Sun, G. et al. (2022). The maize single-nucleus transcriptome comprehensively describes signaling networks governing movement and development of grass stomata. Plant Cell 34: 1890–1911.

Sun, S. et al. (2023). Single-cell RNA sequencing provides a high-resolution roadmap for understanding the multicellular compartmentation of specialized metabolism. Nat. Plants 9: 179–190.

Tian, C., Du, Q., Xu, M., Du, F., and Jiao, Y. (2020). Single-nucleus RNA-seq resolves spatiotemporal developmental trajectories in the tomato shoot apex. bioRxiv: 2020.09.20.305029.

Wang, J.L., Long, J.J., Hotchkiss, T., and Berry, J.O. (1993). C4 photosynthetic gene expression in light- and dark-grown amaranth cotyledons. Plant Physiol. 102: 1085–1093.

Wang, L., Wan, M.C., Liao, R.Y., Xu, J., Xu, Z.G., Xue, H.C., Mai, Y.X., and Wang, J.W. (2023). The maturation and aging trajectory of Marchantia polymorpha at single-cell resolution. Dev. Cell 58: 1429–1444.e6.

Wang, Y., Huan, Q., Li, K., and Qian, W. (2021). Single-cell transcriptome atlas of the leaf and root of rice seedlings. J. Genet. Genomics 48: 881–898.

Weber, E., Engler, C., Gruetzner, R., Werner, S., and Marillonnet, S. (2011). A modular cloning system for standardized assembly of multigene constructs. PLoS One 6: e16765.

Williams, B.P., Burgess, S.J., Reyna-Llorens, I., Knerova, J., Aubry, S., Stanley, S., and Hibberd, J.M. (2016). An Untranslated *cis* -Element Regulates the Accumulation of Multiple C _4_ Enzymes in *Gynandropsis gynandra* Mesophyll Cells. Plant Cell 28: 454–465.

Williams, B.P., Johnston, I.G., Covshoff, S., and Hibberd, J.M. (2013). Phenotypic landscape inference reveals multiple evolutionary paths to C4photosynthesis. Elife 2: 1–19.

Xu, J., Bräutigam, A., Weber, A.P.M., and Zhu, X.G. (2016). Systems analysis of cis-regulatory motifs in C4 photosynthesis genes using maize and rice leaf transcriptomic data during a process of de-etiolation. J. Exp. Bot. 67: 5105–5117.

Zhang, Y., Liu, T., Meyer, C.A., Eeckhoute, J., Johnson, D.S., Bernstein, B.E., Nussbaum, C., Myers, R.M., Brown, M., Li, W., and Shirley, X.S. (2008). Model-based analysis of ChIP-Seq (MACS). Genome Biol. 9.

