## Supplemental Figures for "Single nuclei sequencing reveals C_4_ photosynthesis is based on rewiring of ancestral cell identity networks"

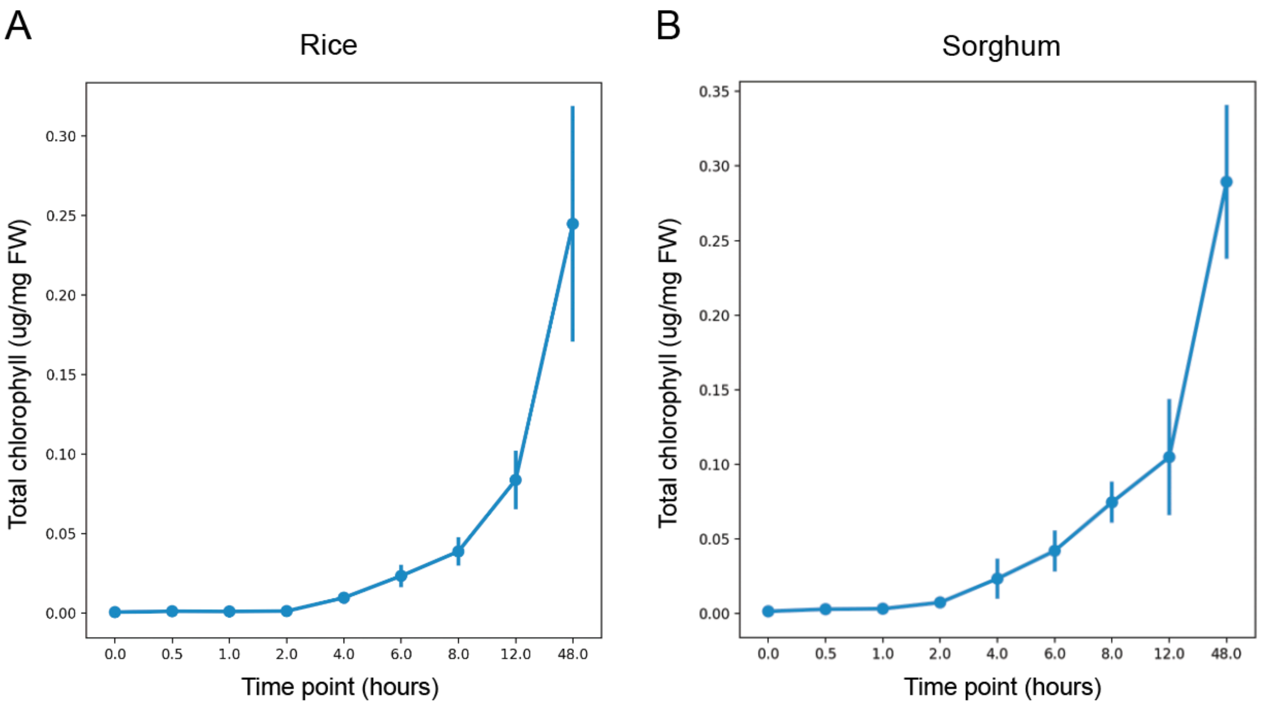


**Figure S1: Chlorophyll accumulation in rice and sorghum seedlings over the time course of de-etiolation.** Total chlorophyll (chlorophyll a + chlorophyll b) measured at different time points during de-etiolation in **(A)** rice and **(B)** sorghum. Each data point represents the mean of 3 biological replicates, +/- standard deviation from the mean.

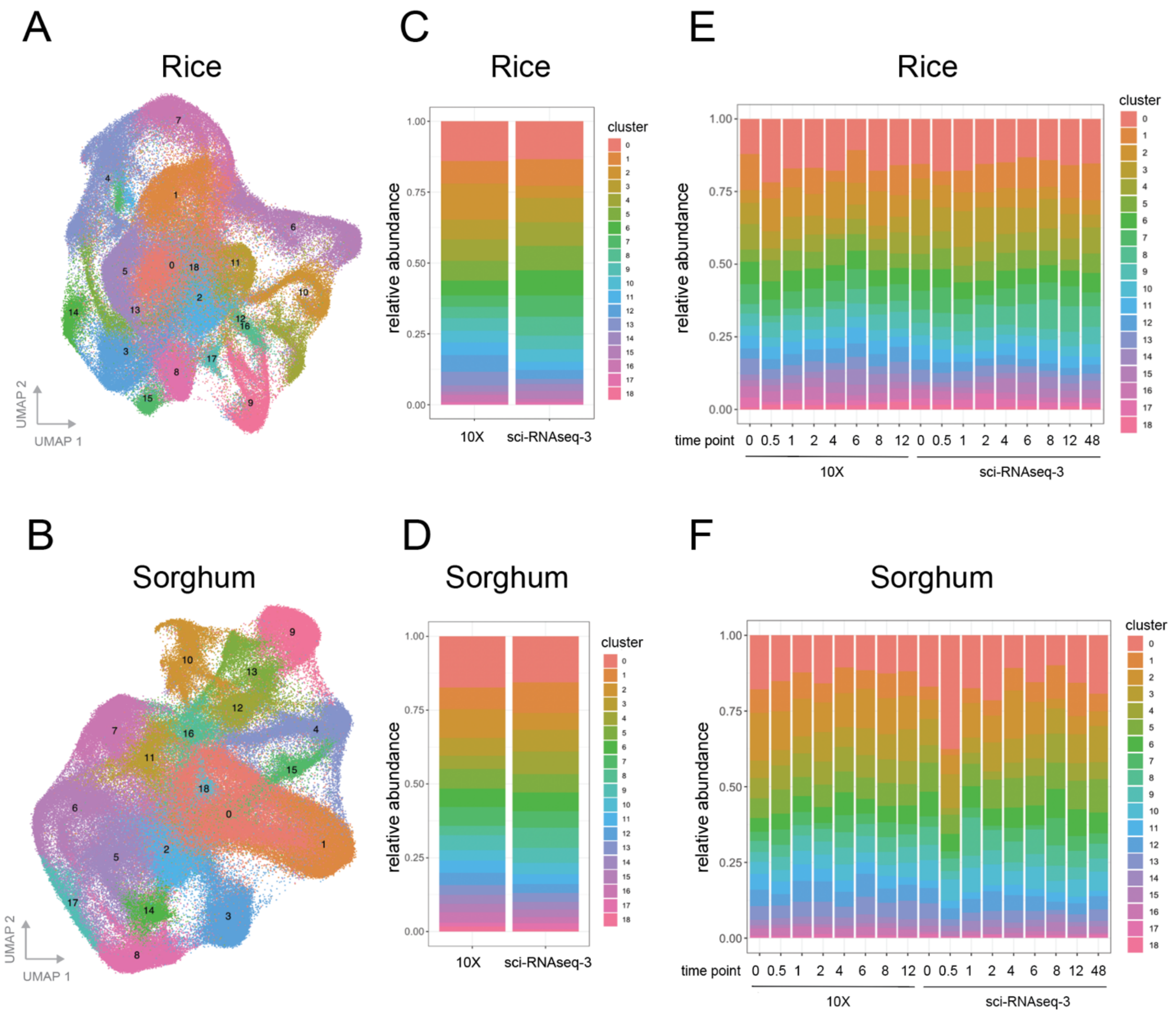


**Figure S2: Single nuclei atlases for gene expression in rice and sorghum shoots during de-etiolation.** Uniform manifold approximation and projection (UMAP) of transcript profiles from single nuclei across **(A)** rice and **(B)** sorghum, across all time points tested. Each cluster in **(C)** rice and **(D)** sorghum contained nuclei sequenced from either by 10X or by sci-RNA-seq3 methods. Similarly, each cluster in **(E)** rice and **(F)** sorghum contained nuclei sequenced from each time point assayed.

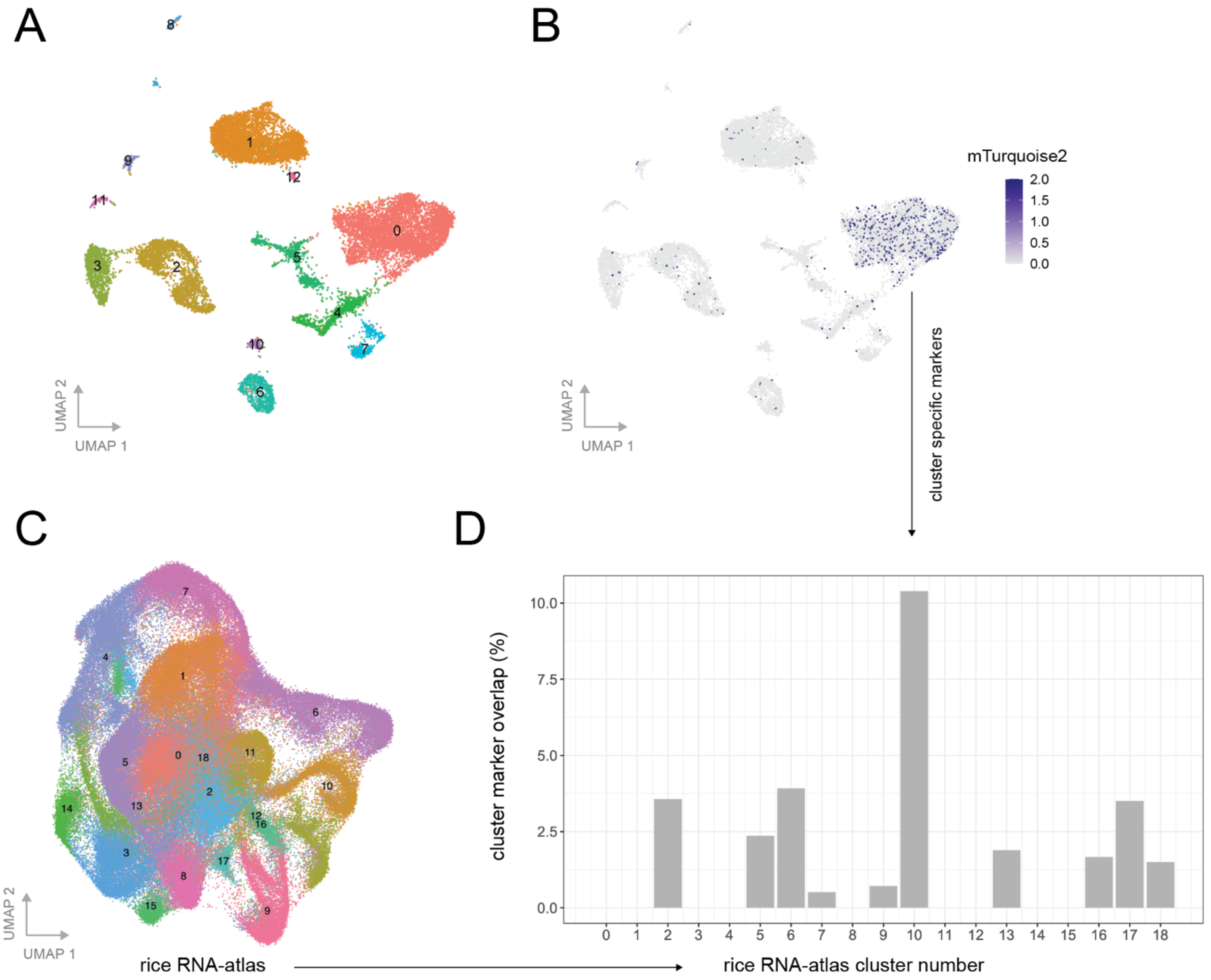


**Figure S3: Bundle sheath transgenic marker line identifies the bundle sheath cluster in the rice transcriptional atlas. (A)** Clustered single nuclei transcript profiles from the rice line expressing mTurquoise2 driven by the bundle sheath specific Zj*PCK* promoter. **(B)** mTurquoise2 expression in the transgenic line. **(C)** Cluster of the rice transcriptional atlas. **(D)** Percent overlap of top cluster markers shared between the bundle sheath transgenic line (B) and the rice transcriptional atlas (C).

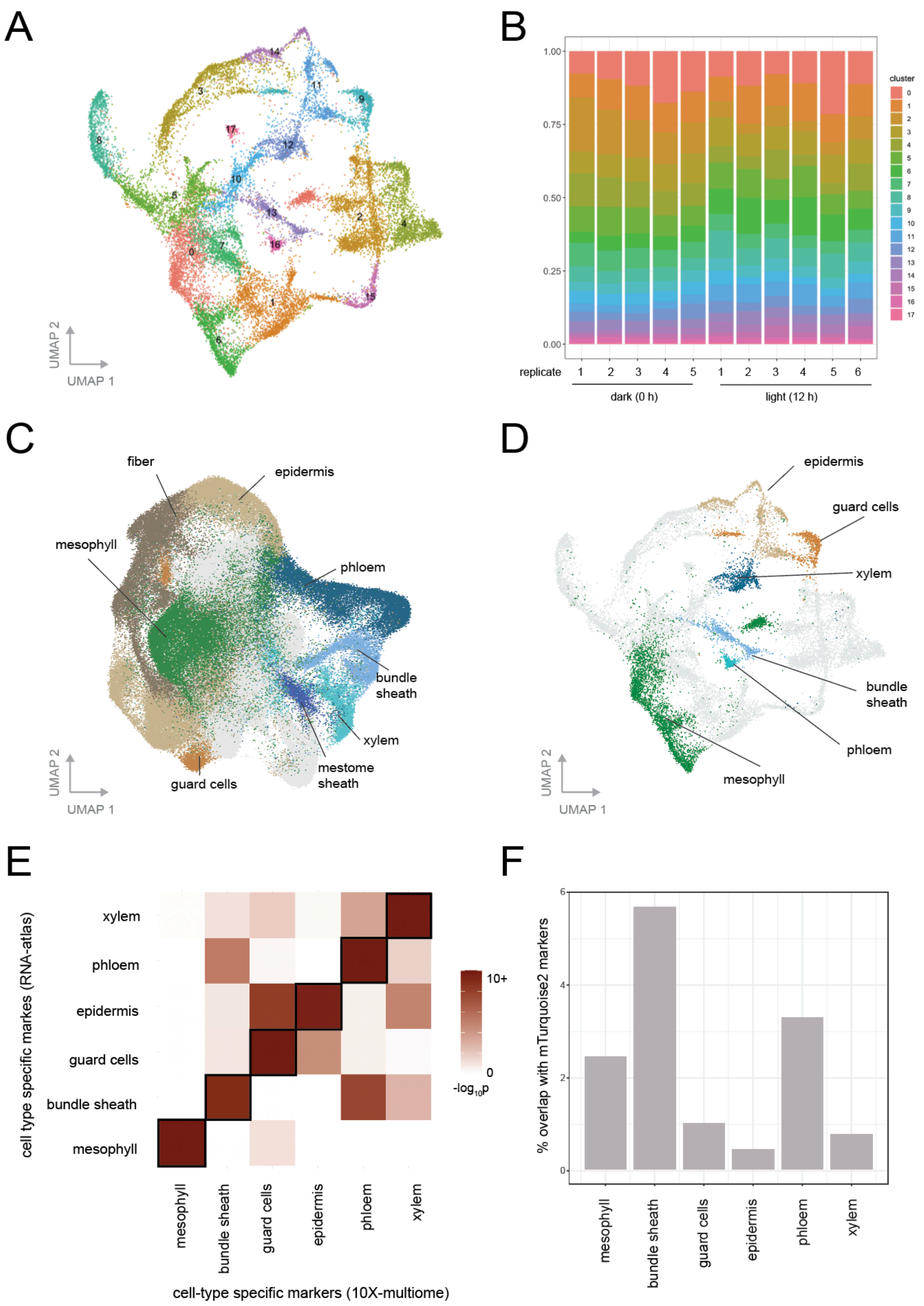


**Figure S4: Identifying cell types in rice 10X-multiome (RNA+ATAC) dataset. (A)** UMAP clustering of 22,154 rice nuclei sequenced using the 10X-multiome workflow. Clustering performed using RNA features. **(B)** Cluster representation of each biological replicate assayed either under etiolated (0h) or light treated (12h) conditions. **(C)** Transcriptional atlas of rice (190,569 nuclei) sequenced using 10X-RNA and sci-RNA-seq3 techniques. **(D)** Transfer of cell transcriptional identities from this main atlas to the 10X-multiome dataset was achieved by **(E)** overlapping significant marker genes from candidate multiome clusters with clusters annotated in the transcriptome dataset. **(F)** Additionally, overlapping the bundle sheath cluster identified in the 10X-multiome data agreed most strongly with markers found in the bundle sheath mTurquoise2 transgenic line.

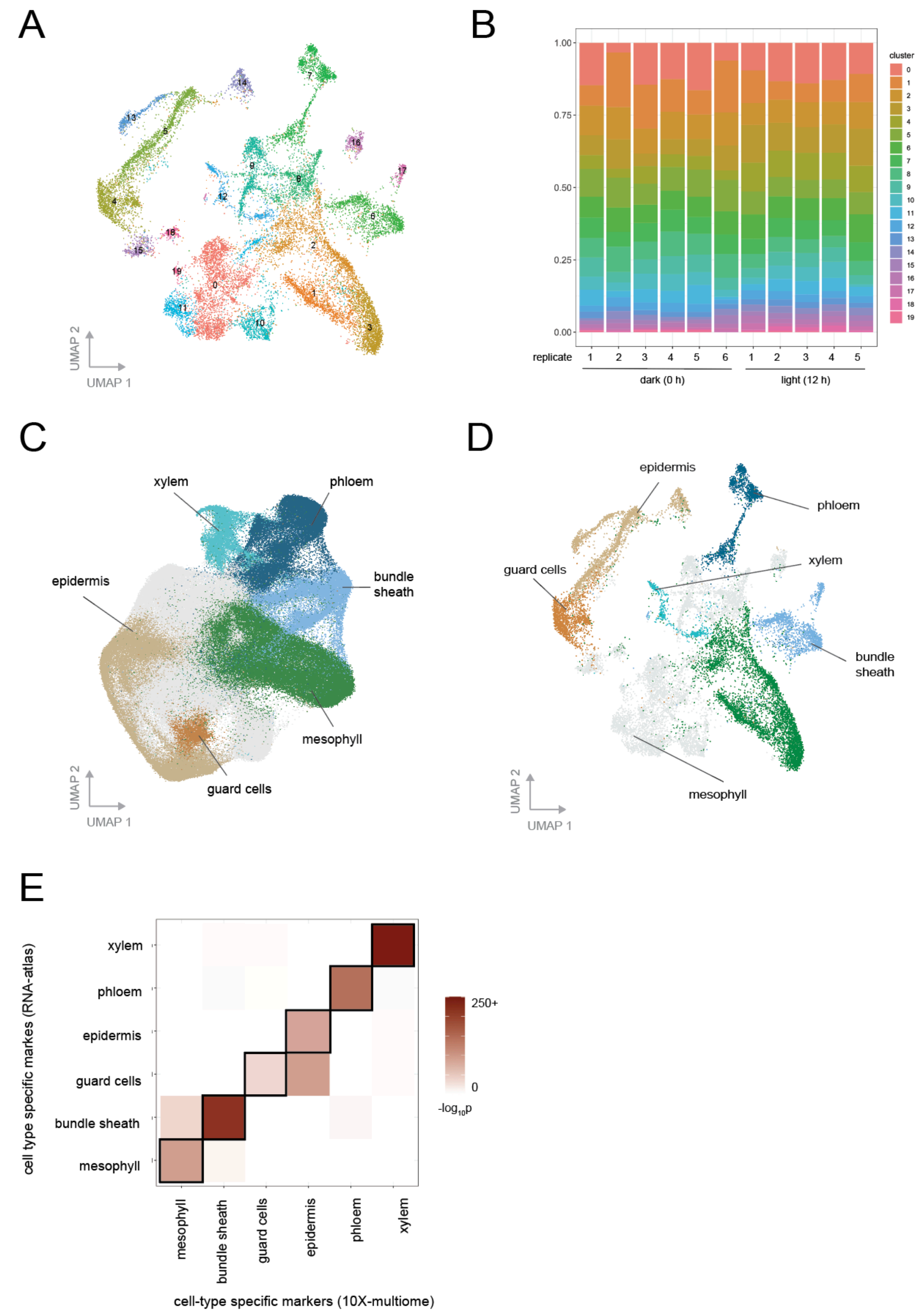


**Figure S5: Identifying cell types in sorghum 10X-multiome (RNA+ATAC) dataset. (A)** UMAP clustering of 20,169 sorghum nuclei sequenced using the 10X-multiome workflow. Clustering was performed using RNA features. **(B)** Cluster representation of each biological replicate assayed either under etiolated (0h) or light treated (12h) conditions. **(C)** Transcriptional atlas of rice (265,701 nuclei) sequenced using 10X-RNA and sci-RNA-seq3 techniques. **(D)** Transfer of cell transcriptional identities from this main atlas to the 10X-multiome dataset was achieved by **(E)** overlapping significant marker genes from candidate multiome clusters with clusters annotated in the transcriptome dataset.

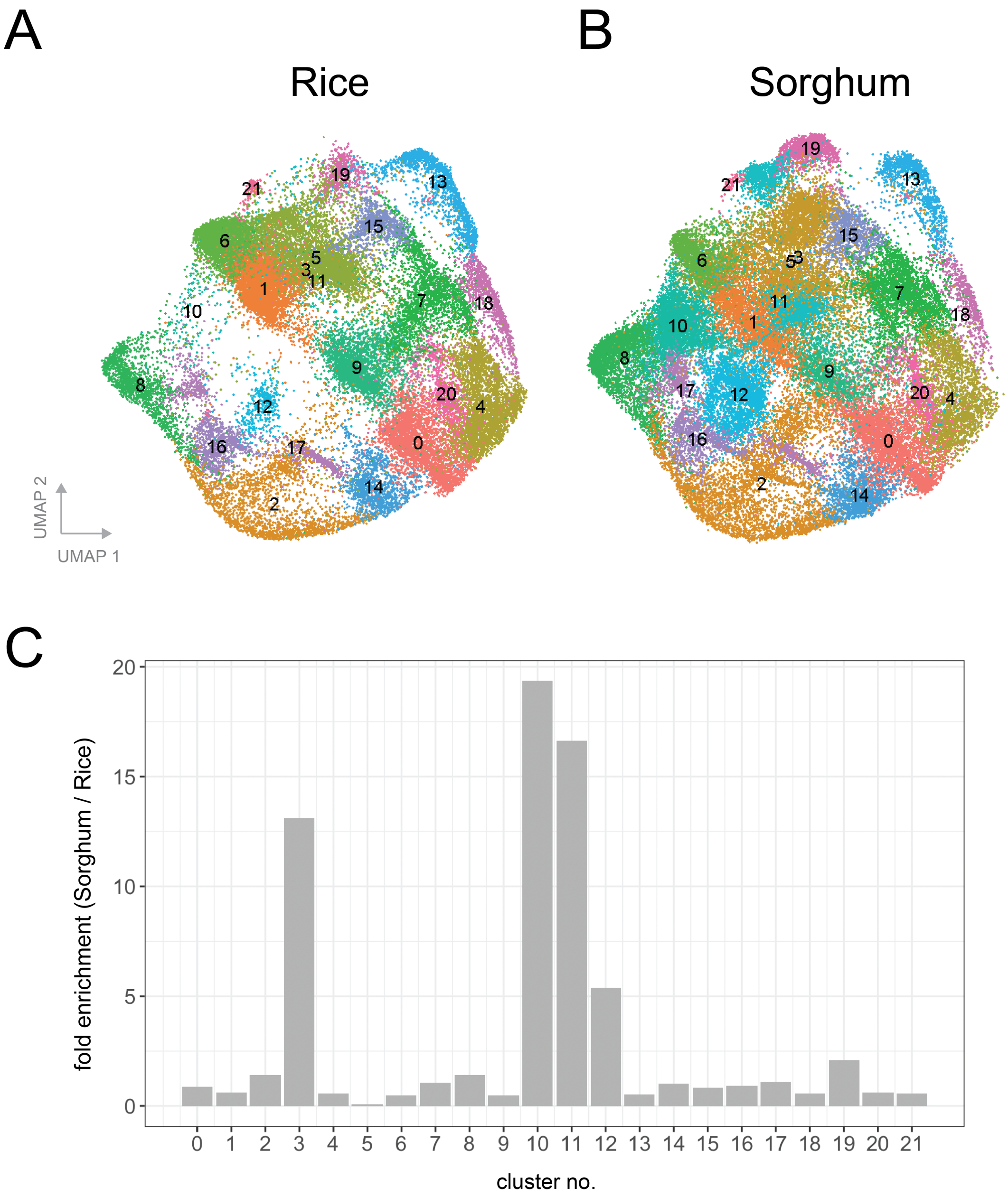


**Figure S6: Clustering of the rice and sorghum pan-transcriptome atlas.** Nuclei clustering visualizing the single nuclei pan-transcriptomes of **(A)** rice and **(B)** sorghum nuclei 48h after light exposure. **(C)** Fold enrichment of cluster membership found in sorghum relative to that of rice.

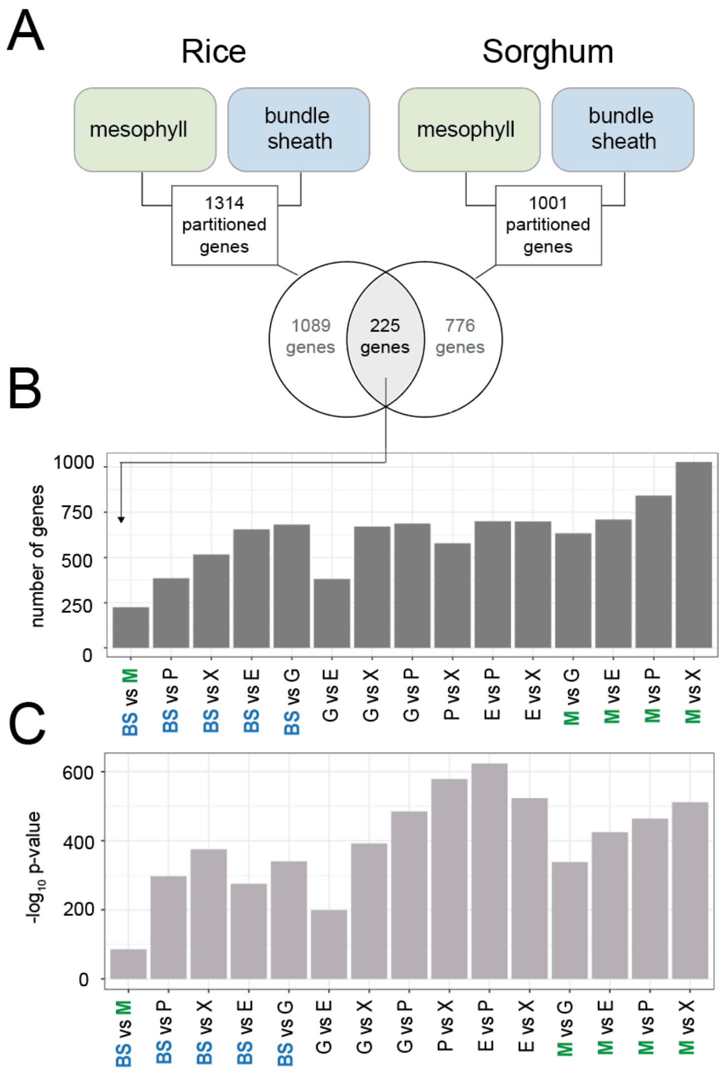


**Figure S7: Bundle sheath and mesophyll cell types show the lowest conservation in transcript partitioning between rice and sorghum. (A)** Number and overlap of genes differentially expressed between mesophyll and bundle sheath cells in rice and sorghum. **(B)** Overlap of genes partitioned between 15 different cell type pairs in rice and sorghum. **(C)** Significance of gene overlap in (B) as found by Fisher Exact Test.

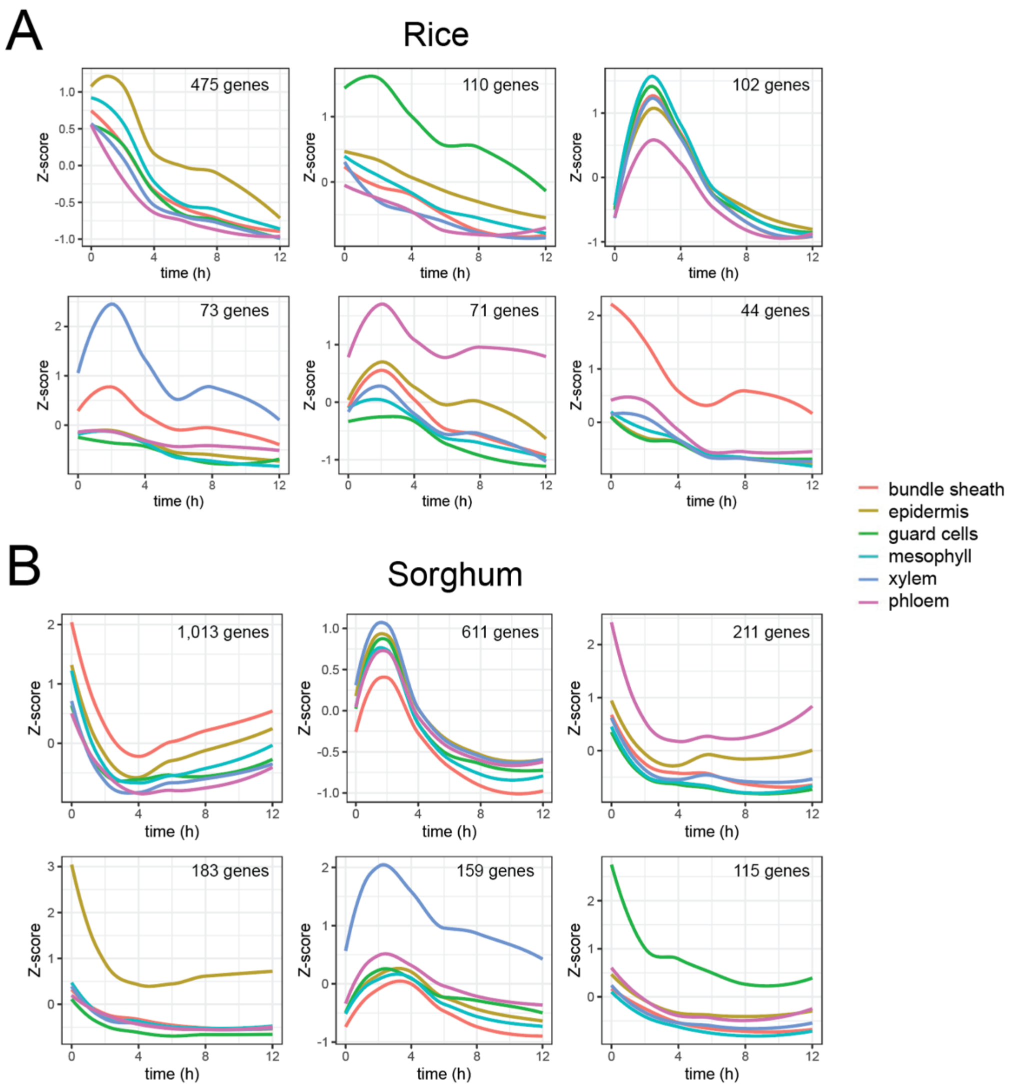


**Figure S8: Light induces changes in cell-type specific transcript abundance.** Repression of transcript abundance from light responsive genes in the first 12h of exposure to light. Clusters of genes were identified using Pearson correlation. Each cluster shows patterns of gene expression induction unique to each cell type in **(A)** rice and **(B)** sorghum.

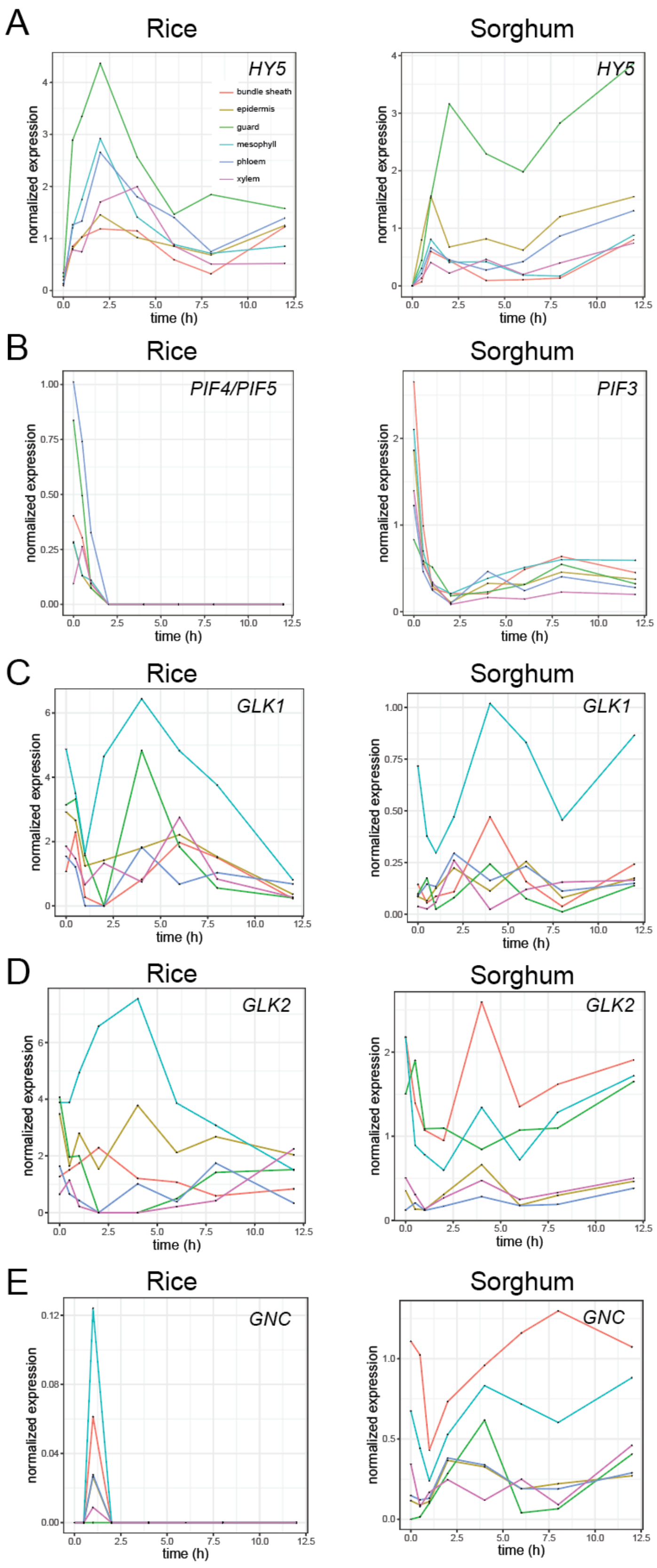


**Figure S9: Cell type specific gene expression patterns of key light transcriptional regulators over time during de-etiolation. (A)** Hy5 expression in rice and sorghum during the first 12h of light exposure across 6 cell types. Similarly, **(B)** PIF, **(C)** GLK1, **(D)** GLK2 and **(E)** GNC expression across each species.

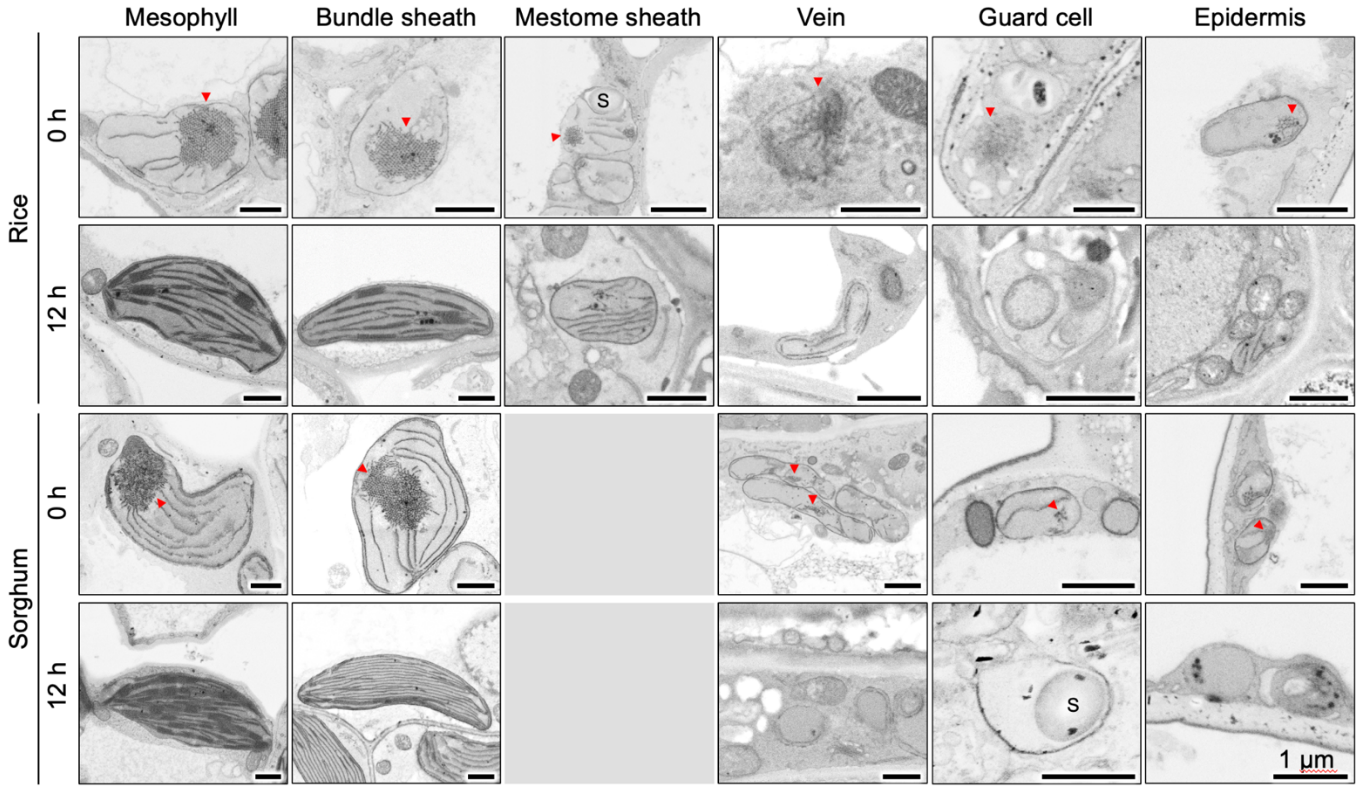


**Figure S10: Scanning electron micrographs of etioplasts and chloroplasts at 0h and 12h after exposure to light in different cell types of rice and sorghum shoots.** Etioplasts are indicated with a red arrowhead. ‘S’ indicates starch granules. Unlike rice, sorghum does not have a mestome sheath.

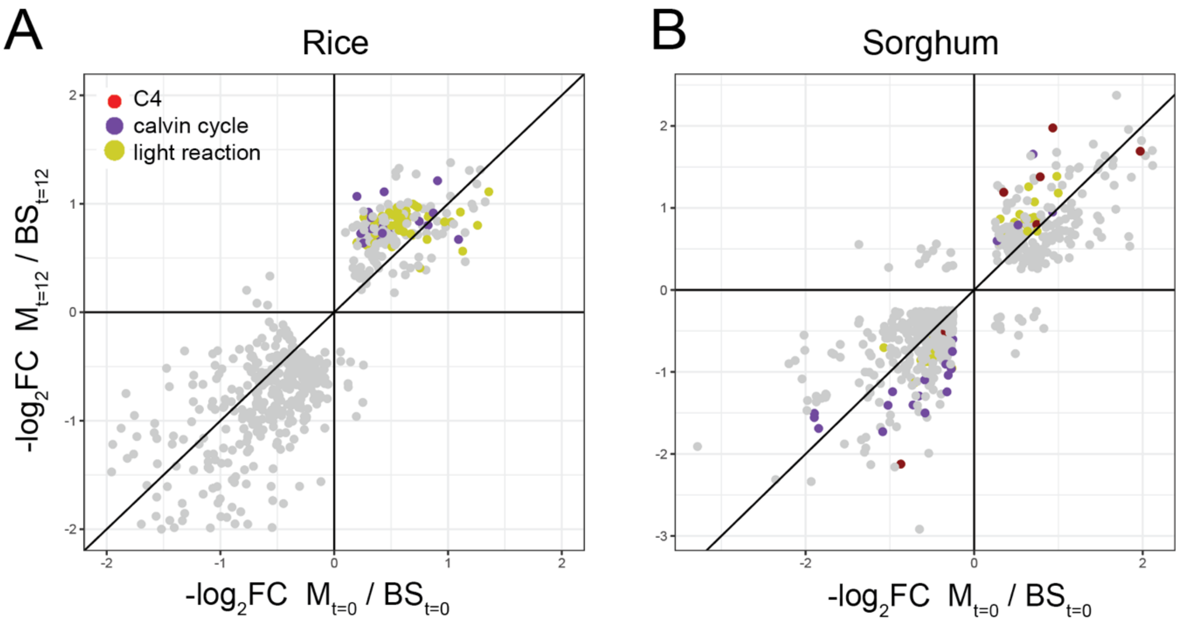


**Figure S11: Light partitions photosynthetic gene expression.** Differences in fold change gene expression in **(A)** rice or **(B)** sorghum genes in the etiolated state (t=0) vs their expression after 12h of light exposure between mesophyll and bundle sheath. Genes encoding enzymes involved in $C_4$ photosynthesis, the Calvin Benson Bassham cycle and the light reactions are shown in red, purple, and yellow, respectively.

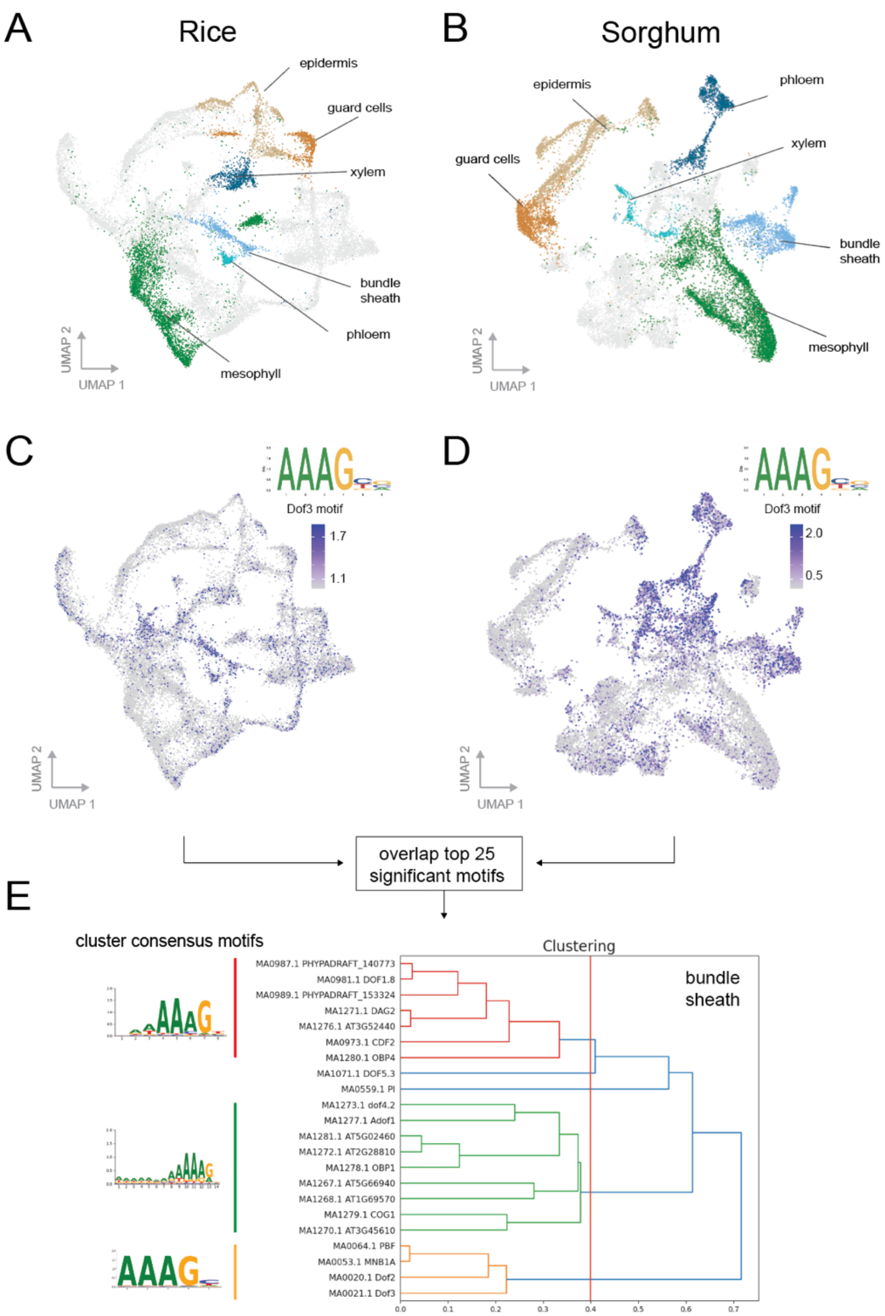


**Figure S12: Discovering conserved cell-type specific *cis*-elements across species.** 10X-multiome UMAPs of **(A)** rice and **(B)** sorghum clustered using RNA as features. Dof3 motif prevalence within accessible chromatin is restricted to bundle sheath and phloem clusters in **(C)** rice and **(D)** sorghum. **(E)** For each species, the top 25 most significantly enriched motifs within the bundle sheath were overlapped, and clustered using Tobias. A threshold of 0.4 was used to find consensus motifs (indicated on left).

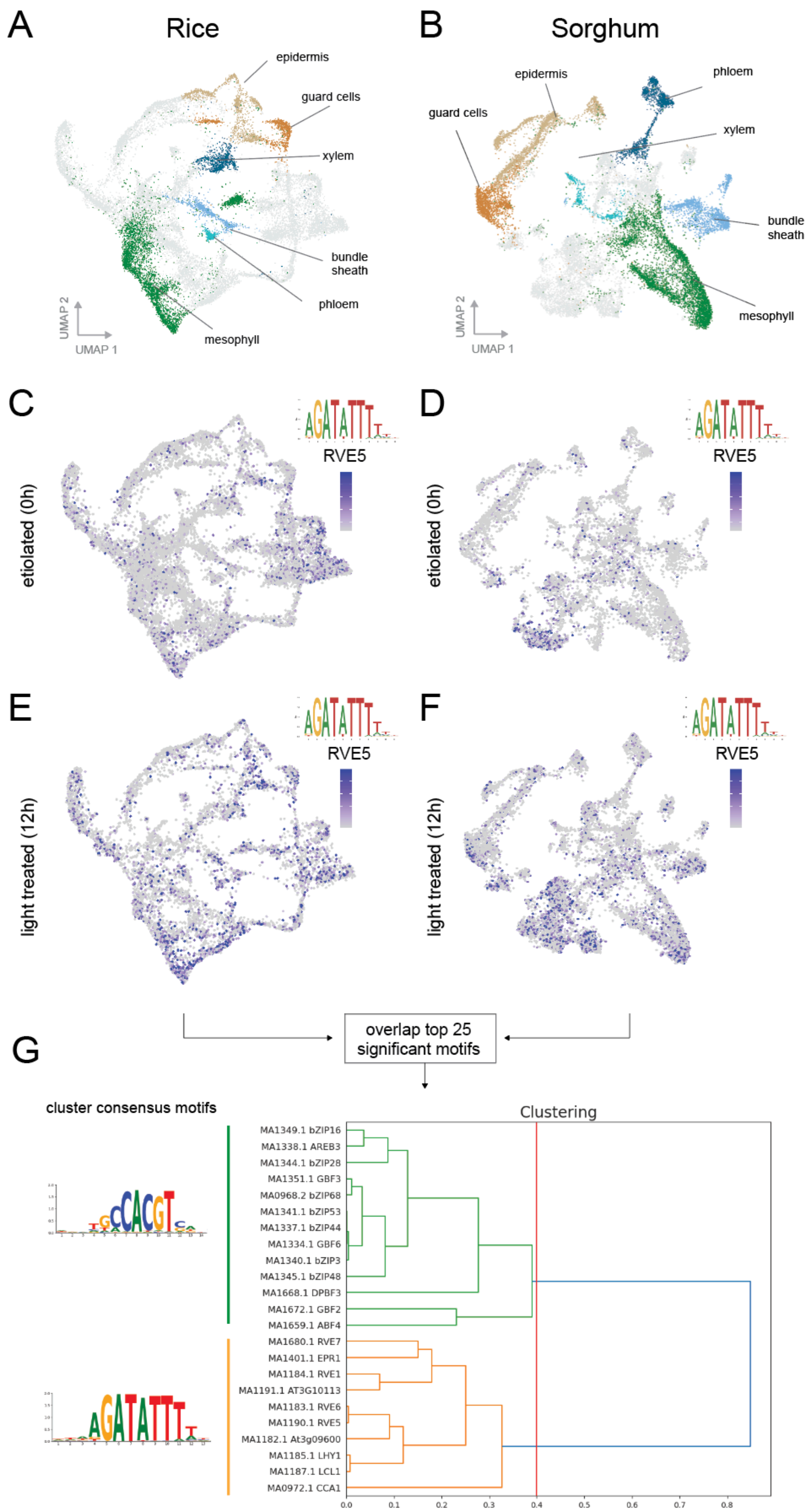


**Figure S13: Discovering conserved light-responsive *cis*-elements across species.** 10X-multiome UMAPs of **(A)** rice and **(B)** sorghum clustered using RNA as features. RVE5 motif prevalence within accessible chromatin is detectable in all cell types in light treated seedlings of **(C&E)** rice and **(D&F)** sorghum. **(G)** For each species, the top 25 most significantly enriched motifs within the bundle sheath were overlapped, and clustered using Tobias. A threshold of 0.4 was used to find consensus motifs.

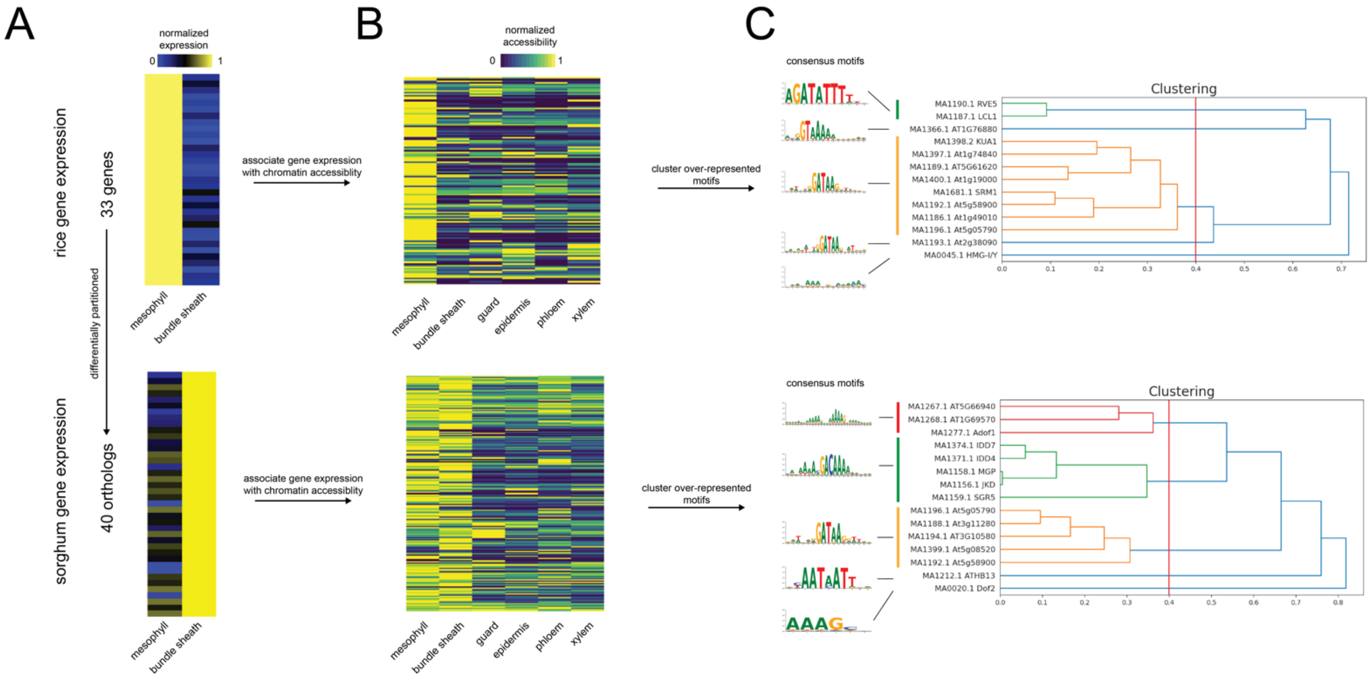


**Figure S14: Discovering *cis*-elements underlying differentially partitioned genes. (A)** Gene expression heatmaps of differentially partitioned genes in rice and sorghum 10X-multiome data, and **(B)** the accessible chromatin that is associated with these expression patterns. **(C)** Enriched motifs were searched for among accessible chromatin, and these motifs were clustered using Tobias to find conserved motifs. Consensus motifs of each cluster are indicated on the left.

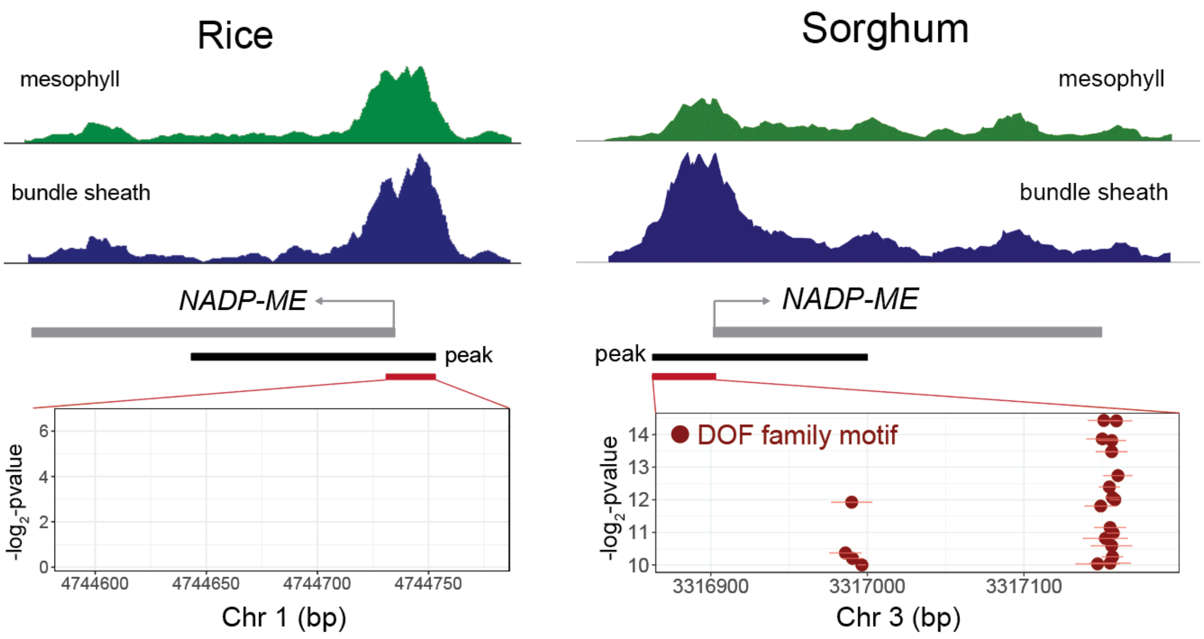


**Figure S15: Mapping accessible chromatin and quantifying DOF motifs in *NADP-ME* promoter.** Chromatin accessibility between mesophyll (green) and bundle sheath (blue) cell types for rice and sorghum for *NADP-ME.* Sequences within accessible chromatin (but not with gene bodies) were analyzed for the presence of DOF family motifs.
